# Dynamic Expansions and Retinal Expression of Spectrally Distinct Short-Wavelength Opsin Genes in Sea Snakes

**DOI:** 10.1101/2024.07.03.602000

**Authors:** Isaac H. Rossetto, Alastair J. Ludington, Bruno F. Simões, Nguyen Van Cao, Kate L. Sanders

## Abstract

The photopigment-encoding visual opsin genes that mediate colour perception show great variation in copy number and adaptive function across vertebrates. An open question is how this variation has been shaped by the interaction of lineage-specific structural genomic architecture and ecological selection pressures. We contribute to this issue by investigating the expansion dynamics and expression of the duplicated Short-Wavelength-Sensitive-1 opsin (SWS1) in sea snakes (Elapidae). We generated one new genome, 45 resequencing datasets, 10 retinal transcriptomes, and 81 SWS1 exon sequences for sea snakes, and analysed these alongside 16 existing genomes for sea snakes and their terrestrial relatives. Our analyses revealed multiple independent transitions in SWS1 copy number in the marine *Hydrophis* clade, with at least three lineages having multiple intact SWS1 genes: the previously studied *Hydrophis cyanocinctus* and at least two close relatives of this species; *H. atriceps*-*H. fasciatus;* and an individual *H. curtus*. In each lineage, gene copy divergence at a key spectral tuning site resulted in distinct UV and Violet/Blue-sensitive SWS1 subtypes. Both spectral variants were simultaneously expressed in the retinae of *H. cyanocinctus* and *H. atriceps,* providing the first evidence that these SWS1 expansions confer novel phenotypes. Finally, chromosome annotation for nine species revealed shared structural features in proximity to SWS1 regardless of copy number. If these features are associated with SWS1 duplication, expanded opsin complements could be more common in snakes than is currently recognised. Alternatively, selection pressures specific to aquatic environments could favour improved chromatic distinction in just some lineages.

**Significance:** Secondary transitions to marine environments are commonly accompanied by pseudogenisation of the visual opsin genes which mediate colour perception. Conversely, a species of fully-marine hydrophiid snake has functionally expanded its short-wavelength-sensitive opsin repertoire following a terrestrial ancestry. The current study explores this further by mapping opsin copy number across the hydrophiid phylogeny and by quantifying expression of SWS1 subtypes within sea snake retinae. Despite few reports of opsin expansions in tetrapods, we provide evidence for the occurrence of multiple expansion events throughout *Hydrophis*. Most intriguingly, retinal expression of spectrally-divergent copies implies a functionally-significant phenotype; possibly even trichromacy.

## Introduction

The remarkable diversity of animal visual systems is underpinned by variation in the photopigment-encoding visual opsin genes that mediate colour perception (Cronin et al. 2014). The ancestor of all vertebrates possessed four spectrally distinct pigments expressed in cone photoreceptors: a Long-Wavelength Opsin (LWS, with maximum wavelength absorbance [λ_max_] of 510 -570 nm), Short-Wavelength Opsin 1 (SWS1; λ_max_ ≈ 358 - 440 nm), Short-Wavelength Opsin 2 (SWS2; λ_max_ ≈ 400-460 nm), and Rhodopsin 2 (RH2; λ_max_ ≈ 470 - 510 nm; Hagen et al. 2023 and references therein). Over the subsequent 400 million years of vertebrate evolution, this ancestral complement of cone opsins has undergone substantial diversification via gene losses, duplications and functional divergence (Hagen et al. 2023). However, while numerous opsin losses have been reported in all major vertebrate groups, these taxa exhibit striking disparity in their propensity for expansions via gene duplication and subsequent functional divergence (Hagen et al. 2023 and references therein). In particular, teleost fish have undergone extensive expansions in multiple independent lineages (Musilova et al. 2021), resulting in complements of up to 38 intact visual opsins (Musilova et al. 2019). Tetrapods, in contrast, are so far known to have undergone only four functional expansions, with the addition of between one and three or four intact genes in each lineage (Jacobs et al. 1996; Hunt et al. 1998; Dulai et al. 1999; Hauzman et al. 2021; Rossetto et al. 2023). The primary drivers of variation in gene copy number are structural genomic architecture (Chen et al. 2014), which underpins propensity for duplication events, and ecological selection pressures, which determine the persistence of genes within populations. An important remaining question is how these evolutionary factors have interacted to shape the striking diversity of opsin copy number and adaptive function in vertebrate visual opsins.

While the genomic mechanisms of visual opsin expansions have been well studied in teleost fish (Musilova et al. 2021 and references therein), very little is known about the origin of new opsins in tetrapods. Whole-genome and chromosome duplication events might have produced the visual opsins of the ancestral vertebrate from a single primitive LWS opsin (Nordström et al. 2004; Larhammar et al. 2009; Lagman et al. 2013). The many opsin subtypes observed in fish lineages appear to be generated by retroduplications (Ward et al. 2008; Rennison et al. 2012; Fujiyabu et al. 2019) or additional lineage-specific whole-genome duplications (Meyer & Van de Peer 2005; Lin et al. 2017). Tandem duplications are also commonly reported in fish, although their exact mechanism is uncertain (Rennison et al. 2012; Musilova et al. 2021 and references therein). The duplications of MWS/LWS in primate lineages are suspected to have occurred by unequal crossing over of sister chromatids during recombination (Nathans et al. 1986; Hunt et al. 1998), sometimes promoted by Alu transposons (Dulai et al. 1999). Whole-genome duplications have not been reported in reptiles (Otto & Whitton 2000; Mable 2004) and visual opsin duplication has been described in only two lineages (Hauzman et al. 2021; Rossetto et al. 2023).

In contrast to studies of molecular mechanisms, much research attention has focused on the adaptive processes that have shaped visual opsin complements in vertebrates, both at recent and macroevolutionary timescales (reviewed in Hagen et al. 2023). There are three ways by which retained gene duplicates can be phenotypically influential: (1) the additional copies increase transcriptional dosage, (2) relaxed selective constraint causes subfunctionalization of some or all copies, and (3) certain gene copies provide novel functions (known as neofunctionalization; Pegueroles et al. 2013). A primary mechanism of functional divergence in visual opsins involves shifts in maximal wavelength absorbance (λ_max_) via amino acid changes at so-called spectral tuning sites. These key sites control the shape of the opsin binding pocket and, alongside the type of vitamin-A derived light-absorbing chromophore, determine the wavelength of light that is most likely to initiate the phototransduction cascade (Yokoyama 2002; Yokoyama & Yokoyama 2003; Corbo 2021).

Interestingly, both of the currently known visual opsin expansions in reptiles have occurred in the SWS1 of aquatic snakes (Hauzman et al. 2021; Rossetto et al. 2023). Most snakes possess only two cone-expressed visual opsins (SWS1 and LWS) following a dim-light bottleneck which resulted in losses of SWS2 and RH2 in their earliest ancestors (Simões et al. 2015). A study of freshwater snakes in the *Helicops* genus (Hauzman et al. 2021) reported dual expression of site 86-diverged UV-sensitive and Violet-sensitive SWS1 opsins (λ_max_ ≈ 363-395 nm and λ_max_ ≈ 411-420 nm respectively; determined by MSP). A subsequent study of the genome of the fully marine blue-banded sea snake, *H. cyanocinctus*, revealed at least three intact copies that have different amino acids at the key position for spectral tuning of SWS1; these are phenylalanine (F86), which confers peak sensitivity to UV-light (λ_max_ ≈ 360 nm based on microspectrophotometry [MSP]: Simões et al. 2016 & 2020) and tyrosine (Y86), which confers violet/blue-light sensitivity (λ_max_ ≈ 428 nm based on MSP: Hart et al. 2012; Simões et al. 2020) (Rossetto et al. 2023).

While visual opsin duplications can confer adaptive benefits to visually-reliant organisms, there are several cases of opsin loss following secondarily-aquatic transitions (Fasick et al. 1998; Peichl & Moutairou 1998; Levenson et al. 2006). It is entirely possible that SWS1 duplications in *H. cyanocinctus* (and possibly also *Helicops*) occurred too recently to have been deleted or disrupted by neutral or negative selection. However, predicting the likelihood of retention of these genes is difficult without a reliable timeframe for their origins. Gene subfunctionalization via exon disruption or deletion would mean that multiple copies are required to fulfil the functional role of the ancestral SWS1 opsin. No evidence for this exists in *H. cyanocinctus* as each gene copy appears to be undisrupted without stop codon insertions (Rossetto et al. 2023). Instead, the divergence of gene copies at a spectral tuning site (86) that is under positive selection in sea snakes (Simões et al. 2020) provides strong evidence for neofunctionalization.

Here, we investigate the dynamics of visual opsin expansion in sea snakes by characterising the copy number distribution and structural architecture of SWS1 in the genomes of sea snakes and several terrestrial outgroups. We then quantified retinal transcription of UV-sensitive (F86) and Violet/Blue light-sensitive (Y86) variants in eight sea snake species to assess whether the new SWS1 copies have resulted in a functionally adaptive phenotype. Our results suggest that this rare case of visual opsin diversification has resulted in a novel visual adaptation to the marine photic environment following ancestral regression.

## Results and Discussion

### Genome Assembly of the Olive Sea Snake, *Aipysurus laevis*

All previously published high-quality sea snake genomes belong to the *Hydrophis* genus. The *Aipysurus laevis* genome presented here is therefore a useful addition to the genomic resources now available for snakes. We generated 87.2Gbp of nanopore sequence data for *A. laevis* and assembled it into a primary assembly of 4,371 contigs with a total length of 2.07Gbp and N50 of 27Mbp. The final assembly length is slightly larger than the predicted genome size of ∼1.84Gbp (supplementary fig.S2; supplementary table.S2). Assembly quality was assessed using sequence and gene completeness metrics. The assembly achieved a k-mer completeness score of 93.7% (supplementary fig.S3A; supplementary table.S2), which is a measure of the recovered solid k-mers in the assembly compared to its high-accuracy sequence data, along with a consensus quality (QV) score of QV32.6, surpassing 99.9% base accuracy (supplementary table.S2). Assessing the gene content of the assembly using BUSCO identified 94.8% of all Tetrapoda BUSCO genes as complete, 0.9% as being duplicated with the remaining 4.3% of genes reported as missing or fragmented (supplementary fig.S3B).

### Distribution and Origins of SWS1 Expansions in Sea Snakes

Genome mining of long-read assemblies for eight species (*H. cyanocinctus, H. major, H. elegans, H. ornatus, H. curtus, A. laevis, N. naja* and *N. scutatus*) detected single SWS1 genes in all species other than *H. cyanocinctus* (supplementary table.S3). Both UV-sensitive and violet/blue-shifted genes were observed in *H. cyanocinctus;* the single-copy SWS1 genes were UV-sensitive in *H. major*, *H. elegans*, *N. naja* and *N. scutatus*, violet/blue-shifted in *H. curtus* and *H. ornatus*, and violet-shifted in *A. laevis*; refer to table.S1 for genome accession numbers and table.S3 for gene coordinates. Single copies of the RH1 and LWS visual opsin genes were observed in all genomes.

For samples which lacked reference genomes, gene copy number was estimated by normalising short-read coverage at the SWS1 locus. We validated this technique by estimating normalised coverage for 5,045 complete BUSCO genes in *H. major* in 11 re-sequencing samples (supplementary fig.S4). These marker genes are evolutionarily expected to appear once within elapid taxa, and are consequently expected to report one-fold coverage when normalised. Across the 11 test samples, the average normalised average coverage for the 5,045 BUSCO genes ranged between 1.02 - 1.21x, indicating that our methodology was capable of accurately inferring gene copy number from the normalised coverage value (supplementary fig.S4).

For the eight species with long-read assemblies, SWS1 gene copy number was interpreted alongside short-read data for one to eight individuals per species. In the seven species with single copies of SWS1, 22 out of 23 individuals showed 1-fold normalised coverage (0.50x to 1.17x) across the length of the SWS1 gene region. Because these results provided confidence in copy numbers estimated using short read coverage, we tentatively inferred copy numbers for the 10 additional species for which only short read data were available. The semi-aquatic *Hydrelaps* showed one-fold coverage, indicating the presence of a single SWS1 gene. The marine *A. laevis* also has a single SWS1 (based on long and short read data), hence our results provide evidence of SWS1 duplication exclusively within *Hydrophis*. Within this clade, *H. peronii, H. stokesii, H. ocellatus, H. platurus* and *H. schistosus* show one-fold coverage of SWS1 indicating the presence of a single gene. *H. spiralis* and *H. atriceps* showed at least two-fold coverage at the SWS1 locus relative to the chromosome averages, and thus appear to possess two SWS1 genes each (fig.1).

**Figure.1:**
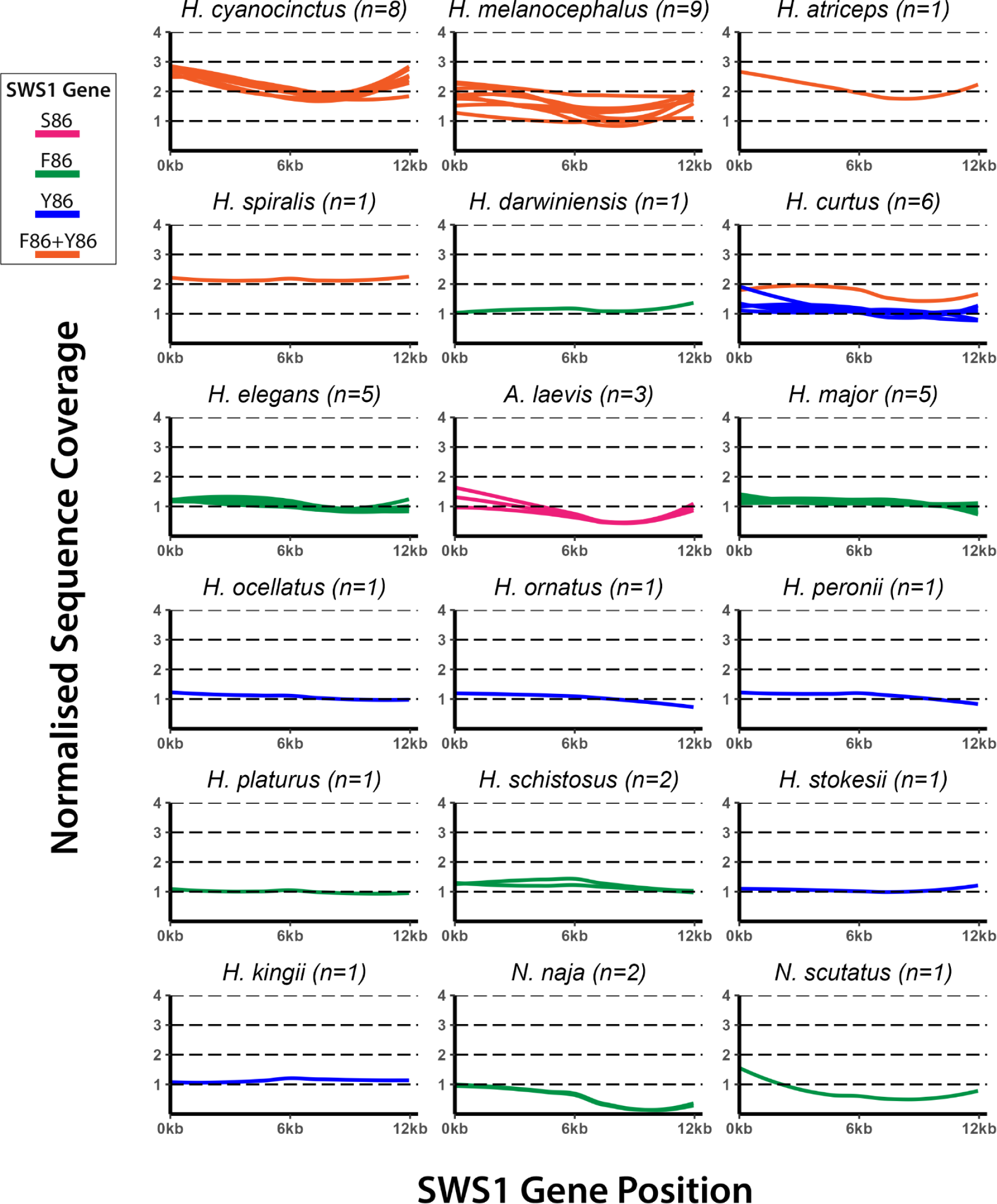
Coverage at SWS1 locus relative to chromosome average for short-read assemblies. See supplementary table.S1 for a full list of samples included. Each data line represents a resequencing sample (sample size included in graph titles). Data is smoothed using loess local regression (span = 1) and grey shading represents bands of confidence. Orange = multiple SWS1 (λmax ≈ 360 nm and 428 nm), Green = single SWS1 (λmax ≈ 360 nm), Blue = single SWS1 (λmax ≈ 428 nm), Pink = single SWS1 (λmax ≈ 420 nm)(as determined by cDNA sequences and from MSP data described in Simões et al. 2016 & 2020).

In *H. cyanocinctus*, the reference assembly contained at least three SWS1 genes, and short-read data for all eight sampled individuals show between two- and three-fold normalised coverage (average range from 2.17x to 2.74x) across the ∼12kb SWS1 region. For these samples, low average sequencing depth resulted in a lower breadth of coverage across chromosome 4 (supplementary table.S7). Accurate estimation of gene copy number by normalisation is dependent on consistent breadth and depth of coverage across the genome, meaning this remains an approximate method for genomes with low depth and no reference genome available. This applies to the short-read data of the nine sampled individuals of *H. melanocephalus* which each vary greatly in depth (2-7x) resulting in variable normalised coverage at the SWS1 site between one and three times the chromosome average. Despite this uncertainty, observations of recent introgression with *H. cyanocinctus* (Sanders et al. 2013) and their recent divergence (<0.5 Mya; Sanders et al. 2008; Ludington et al. 2023) would suggest that *H. melanocephalus* has at least two SWS1 genes (fig.1)

It must be noted that all resequencing data (fig.1) was mapped to *H. major* using the same filter criteria and mapping parameters. However, species which are phylogenetically distant from *H. major* are also likely to be more genetically divergent. Consequently, the average depth and breadth of coverage in these samples will be reduced due to the filtering of accurate but divergent alignments, resulting in a normalised coverage which is lower than expected. This is observed in the coverage profiles of the terrestrial species *N. naja* and *N. scutatus* (fig.1) which diverged from *H. major* some ∼32 Mya and ∼20 Mya, respectively (Lee et al. 2016). The ∼5000 bp intron gulf located in the second half of the SWS1 gene may have diverged more rapidly than the rest of the gene, resulting in reduced normalised coverage than in the exon-rich first half. Despite this, short-read analysis still suggests that *N. naja* and *N. scutatus* each have one SWS1 gene (fig.1), which is confirmed by the genome data (supplementary table.S3).

Relative coverage analysis of one short-read *H. curtus* genome indicated the presence of two SWS1 genes in this individual (fig.1) while a single violet/blue-shifted copy was inferred in the seven other genomes sampled for this species (five short-read and two long-read). Variant calling at site 86 for the individual with two copies revealed that these have also diverged at the spectral tuning site, with one UV-sensitive and one violet/blue-sensitive gene (supplementary table.S4). Although this variation could instead represent polymorphism between alleles, no evidence for heterozygosity at site 86 was observed within the SWS1 genes of single-copy species (supplementary table.S4), suggesting the presence of diverged gene copies. Intriguingly, all six individuals for which short-read data were generated had been sourced from the Gulf of Thailand, indicating that SWS1 copy number may differ within populations of the same species.

Sanger sequencing of SWS1 exon 1 revealed only single peaks at site 86 in all 42 individuals of the 14 species inferred to have single SWS1 genes (using long read assemblies for five species, and short read data for the other nine; data from Simoes et al 2020). In contrast, an excess of double (FY) peaks was found in individuals within populations of species inferred to have multiple SWS1 copies; these were *H. cyanocinctus* (long read), *H. spiralis* (short read) and *H. atriceps* (short read; fig.2). While proportions of alleles based on Hardy–Weinberg expectations provide, at best, indirect evidence of copy number, the results provide support for our inferences from the genomes of these species. Four additional species had double (FY) Sanger peaks but lack whole-genome data. These include all nine sequenced *H. fasciatus* individuals; this species is a close sister lineage to *H. atriceps* (Sanders et al. 2013), which was inferred here to have multiple SWS1 copies using short read data. Similarly, *H. coggeri*, *H. parviceps* and *H. belcheri* have double (FY) Sanger peaks (supplementary table.S1) and are phylogenetically intermediate between *H. cyanocinctus*-*H. melanocephalus* and *H. spiralis*, thus are likely to share multiple SWS1 copies by descent.

**Figure.2:**
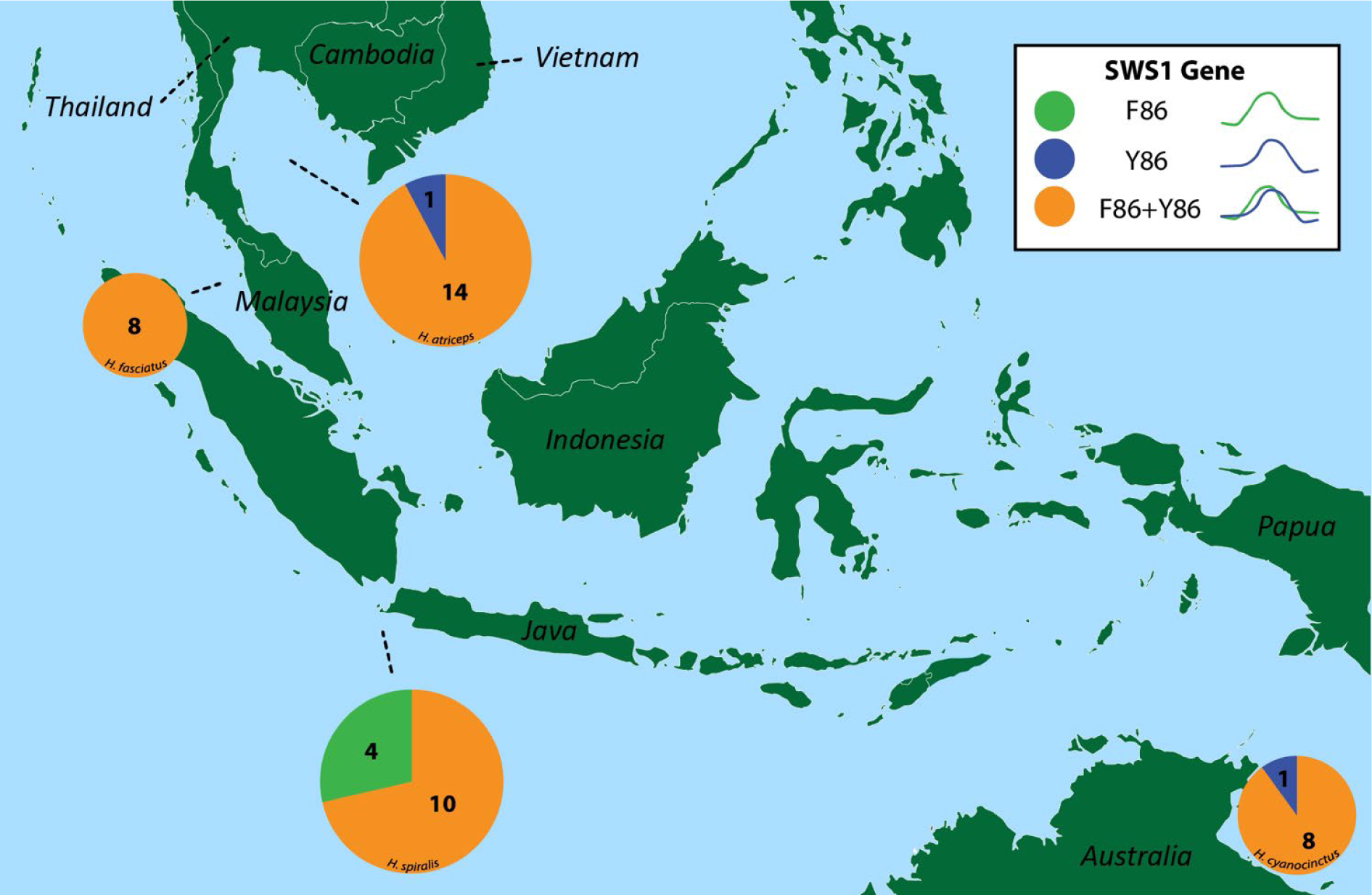
SWS1 site 86 genotype ratios in sea snake populations. To ensure that displayed genotype ratios accurately represent the greater populations, a minimum sample size (n) of 8 individuals per species and locality were sequenced and included in pie charts. Pie chart diameter is relative to sample size, which is also captioned for each pie segment. All data was collected from sanger sequencing of the SWS1 opsin gene exon 1. See supplementary table.S1 for a full list of genotype data.

Sanger sequences with single (F or Y) peaks were observed in some individuals for species confirmed to have multiple SWS1 genes by long-read genome or coverage analysis (fig.2). PCR amplification bias is a possible explanation for this. There are no SNPs between copies at the primer binding sites, though copy-specific secondary structures and polymerase preferences can influence binding efficiency and thus facilitate bias (Pan et al. 2014). Nucleotide polymorphisms are also possible at spectral tuning site 86, though variant calling showed no evidence for this in genomes with single SWS1 copies (supplementary table.S3). Alternatively, SWS1 copy number could indeed vary within species, as appears to be the case in *H. curtus* (fig.1).

These results suggest that, despite the scarcity of reported tetrapod visual opsin expansions, multiple independent duplications of SWS1 may have occurred in sea snakes within the last ∼2.5 million years (fig.3). Under this scenario, the first duplication occurred in the lineage leading to *H. curtus*, closely followed by a convergent duplication in the common ancestor of *H. cyanocinctus*-*H. melanocephalus* and *H. spiralis* and then by a further duplication in the lineage leading to *H. cyanocinctus*. An additional, convergent duplication then occurred in the most recent common ancestor of *H. atriceps* and *H. fasciatus*. Also plausible is a scenario under which a single expansion event occurred ∼3 Mya in the progenitor of *Hydrophis* and was followed by independent gene losses in five lineages (and retention in four) and a later duplication in *H. cyanocinctus* alone (fig.3, left). While we are unable to resolve the pattern of SWS1 gains and losses in this system, it is notable that no traces of SWS1 pseudogenes were detected in the long read genomes of *H. elegans*, *H. ornatus* and *H. major*. Under the model of a single expansion, *H. elegans* and the clade including *H. ornatus* and *H. major* would have independently undergone losses of SWS1 within the last ∼2 million years. What is clear is that this gene has undergone highly dynamic and rapid evolution in *Hydrophis* sea snakes.

**Figure.3:**
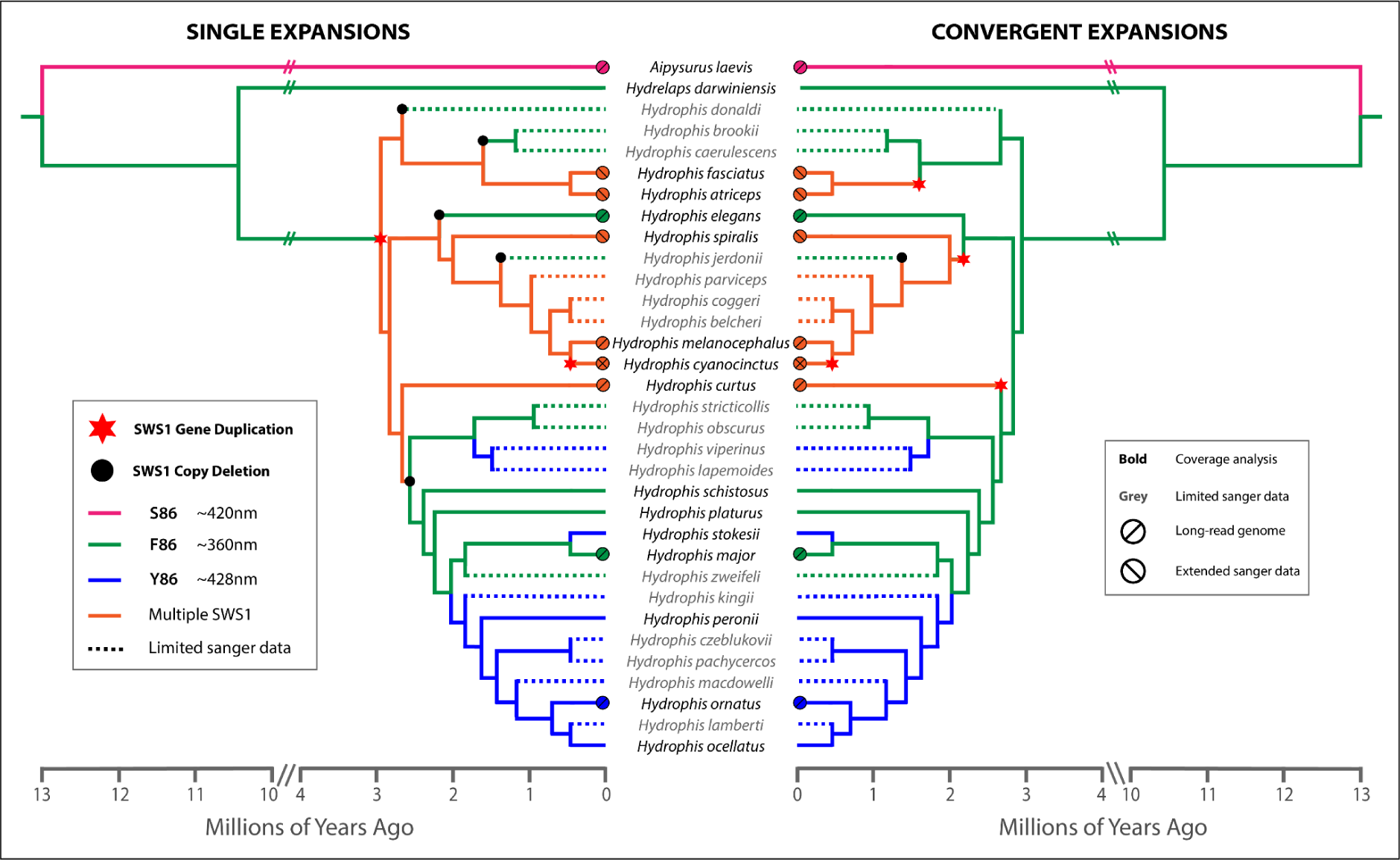
Possible phylogenetic origins of the SWS1 expansion in sea snakes. The ‘Single Event’ phylogeny depicts two duplication events followed by multiple independent gene copy losses. The ‘Convergent Events’ phylogeny depicts four independent duplication events followed by gene copy loss in a single lineage. Phylogenetic data is adapted from the Hydrophis tree presented in Ludington et al. (2023) with Aipysurus and Hydrelaps lineages inferred from the phylogeny presented in Sanders et al. (2008). Visual pigment peak absorbance (λmax) values were predicted from cDNA sequences and from MSP data described in Simões et al. (2020). The lineages labelled with ‘Multiple SWS1’ indicate the likely presence of at least one SWS1-F86 copy and one SWS1-Y86 copy. **‘Coverage analysis’** = species for which coverage analysis has provided strong evidence of single/duplicated genes as depicted in fig.1. **‘Limited sanger data’** = species for which limited SWS1 exon 1 sanger sequencing evidence is available (see supplementary table.S1 for full sample details). **‘Long-read genome’** = species for which long-read genome assemblies provide strong evidence for single/duplicated genes (see supplementary table.S3). **‘Extended sanger data’** = species for which extensive SWS1 exon 1 sanger sequencing data (n=>8 per population) provides evidence for gene duplication as depicted in fig.2.

### Molecular Origin of SWS1 Expansion

Copy number variation is not randomly distributed in genomes. Annotation of chromosomes containing the SWS1 gene for the ten species for which long read genomes were available provided insights into the possible role of genomic architecture in these opsin expansion events (fig.4, supplementary figs.S6-11, table.S5). In *H. cyanocinctus*, all four genes are nested within near-identical ∼46kb repeat segments (fig.4i). Copy D is presumably the ancestral gene due to its chromosomal homology with the single SWS1 in *H. curtus* (Rossetto et al. 2023). However, further comparative synteny analyses were not possible because the genomic region surrounding the repeat segments containing copies A-C does not share homology with the other snakes. Repeat analysis revealed no transposable elements capable of large-scale sequence duplication at repeat boundaries in *H. cyanocinctus*, or at equivalent sites in *H. major* and *H. curtus*. This case of visual opsin expansion is therefore unlikely to have been facilitated by transpositional activity.

**Figure.4:**
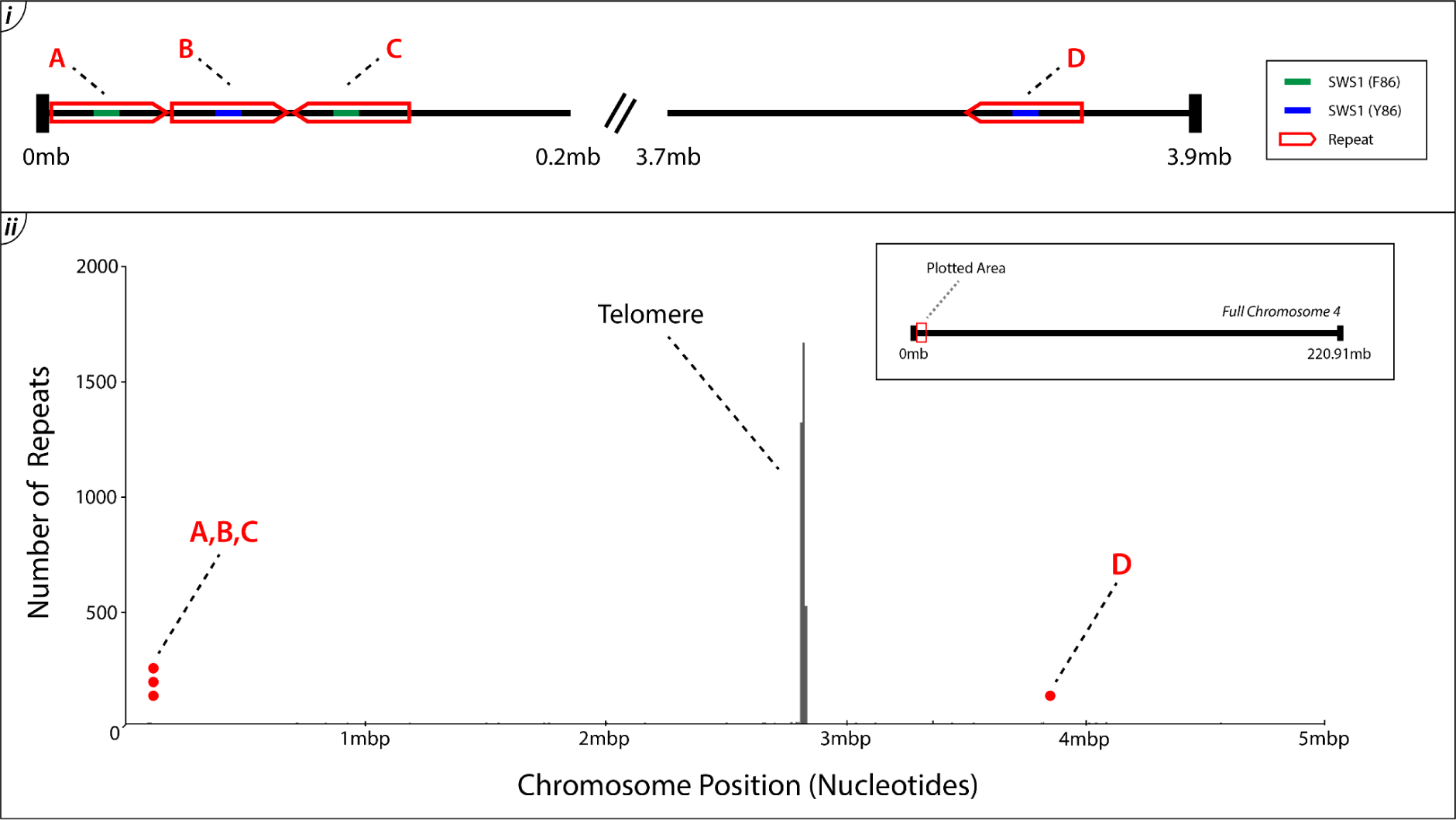
i) Duplicated regions encompassing the SWS1 genes in H. cyanocinctus. ii) Telomere proximity to SWS1 genes on Chromosome 4 of the H. cyanocinctus genome. SWS1 gene locations on the H. cyanocinctus genome (Li et al. 2021; Genbank = GCA_019473425.1) were taken from Rossetto et al. 2023. The y-axis displays the number of TTAGGG telomere repeat sequences present at a particular chromosome position on the x-axis.

Subtelomeric regions are inherently unstable due to high rates of recombination and telomere shortening, rendering them highly susceptible to genomic rearrangements, sequence duplications and deletions (Horvath et al. 2001; Carlton et al. 2002; Mefford & Trask 2002; Bailey et al. 2002; Eichler & Sankoff 2003; See et al. 2006; Shay and Wright 2019). Tandem gene duplications can occur during chromosomal recombination if sister chromatids are misaligned prior to the exchange of genetic material (Reams & Roth 2015). SWS1 genes were observed in close proximity to telomeric repeat sequences in all sea snake genomes (including all copies in *H. cyanocinctus* [fig.4ii]) and in most terrestrial snake genomes (supplementary figs.S6-11, supplementary table.S5). Furthermore, the tail-head orientation of repeated sequences, which is observed in the copy A/B tandem repeat, is often a sign of duplication following misalignment (Nathans et al. 1986; fig.4i). Another potential cause of gene duplication is polymerase slippage during DNA replication (Chen et al. 2005). Twelve strings of 8-26 repeated adenines or thymines (depending on copy orientation) were identified downstream of each SWS1 gene in *Hydrophis*. Any of these repeats could cause polymerase to switch strands during DNA replication, thereby creating an inverted duplicate of the preceding region before reassociating with the original strand. The tail-tail orientation of the copy B/C inverted tandem repeat (a palindrome) might suggest such an occurrence. Moreover, inverted duplicate sequences are known to propagate further duplication (Passananti et al. 1987). This may have facilitated the additional duplication exclusive to *H. cyanocinctus*, following the initial event(s) which produced the two copies observed in other *Hydrophis*.

These results suggest a possible role for genomic features in the expansion of SWS1 genes in *Hydrophis*. However, given that some features are shared by *Hydrophis* species with single as well as multiple gene copies, and the more distantly related *Aipysurus* and terrestrial snakes (fig.4; supplementary figs.S6-11; table.S5), structural architecture alone might not account for SWS1 copy number variation in sea snakes and their terrestrial relatives.

### Retinal Expression and Adaptive Significance of SWS1 Copies

As expected, single SWS1 copies were expressed in the retinae of *A. laevis*, *A. mosaicus*, *H. peronii*, *H. kingii*, *H. darwiniensis* and *H. major* (fig.5), all of which were inferred to have a single copy of SWS1 based on genomic analysis (fig.1-3, supplementary table.S3, Simões et al. 2020). In contrast, multiple spectrally-divergent SWS1 genes were expressed in retinae of *H. cyanocinctus* and *H. atriceps* (fig.5). The high degree of exon homology meant that we could not attribute transcripts to specific copies. However, expression of spectrally-divergent opsins suggests that their expansion might confer a functional phenotype. Trichromacy may be achieved via the simultaneous activation of the three cone-type photopigments found in SWS-duplicated *Hydrophis*: LWS opsin (λ_max_ ≈ 560 nm; Simões et al. 2020), UV-sensitive SWS1 opsin (λ_max_ ≈ 360 nm; Simões et al. 2016 & 2020) and Violet/Blue-sensitive SWS1 opsin (λ_max_ ≈ 428 nm; Hart et al. 2012; Simões et al. 2020). Activated cone photoreceptor cells will either co-express both SWS1 spectral variants simultaneously or express one exclusively. The latter would result in the independent excitation of SWS1 subtypes and facilitate the signal opponency required for trichromacy. The mutually exclusive expression of MWS and LWS opsins in humans is coordinated by an upstream locus control region (LCR) which will randomly loop to the promoter of either opsin gene (Smallwood et al. 2002). This is made possible by their tandem organization, a feature shared by SWS1 copies A-C in *H. cyanocinctus* (fig.4i). Neuroanatomical and behavioural studies are ultimately needed to determine the capacity for trichromacy in SWS-duplicated snakes.

**Figure.5:**
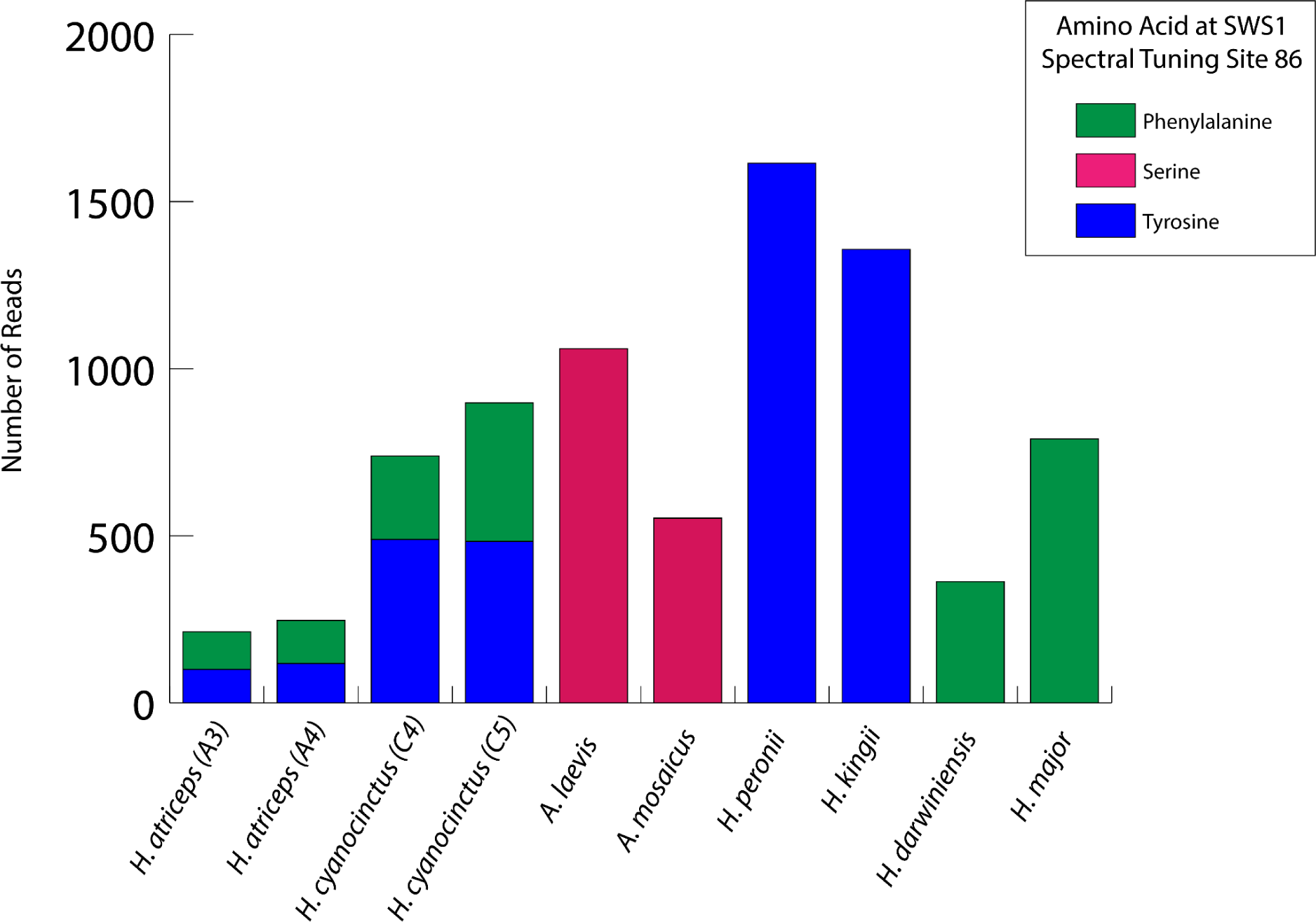
Transcriptional expression at a key spectral tuning site within the SWS1 gene of 10 elapid snakes. The site of interest is spectral tuning site 86 within exon 1 of the SWS1 gene. Bar plots display total transcript counts of each genotype depicted in the legend. Refer to supplementary table.S6 for complete read count data.

While opsin expression patterns have been scarcely investigated in secondarily-aquatic tetrapods, this is not the case for teleost fish (Archer et al. 1997; Britt et al. 2001; Carleton et al. 2008; Seehausen et al. 2008; Dalton et al. 2014; Stieb et al. 2016; Luehrmann et al. 2018 etc.). Interestingly, expression of Short-Wavelength opsin classes (SWS1 λ_max_ ≈ 370 nm and SWS2B λ_max_ ≈ 408 nm) is not static in certain reef fish (Luehrmann et al. 2018). Expression is instead adjusted in response to seasonal and depth-related changes in environmental spectra (Luehrmann et al. 2018). Fully-marine *Hydrophis* snakes may utilize a similar mechanism for differential expression of SWS1 subtypes under variable photic regimes. This flexibility would benefit small-eyed species with limited photoreceptor cells available for opsin expression. Many SWS-duplicated species (*H. atriceps*, *H. fasciatus*, *H. parviceps*, *H. coggeri*, *H. melanocephalus,* and possibly the most recent common ancestor of *H. spiralis* and *H. cyanocinctus*) have evolved tiny heads to specialise on burrowing eel prey, (Sherratt et al. 2018) and consequently have reduced eye size. The topology of photoreceptor cells may also correspond with the underwater visual field, as is reported in some teleosts (Dalton et al. 2014). For example, regional opsin expression could match the incoming light wavelengths of distinct backgrounds if UV-attuned opsins are expressed in ventral photoreceptor populations and blue-attuned opsins are expressed in dorsal populations. This would maximise perception of the photic environment, enabling snakes to better detect predators, prey and potential mates throughout the water column. Single-cell transcriptomics, long-read sequencing of full-length transcripts, or RNA-FISH are required to determine these complex patterns of expression; although the high exon homology between gene copies (>99%) may not permit such analyses.

Hauzman et al. (2021) reported dual expression of site 86-diverged SWS1 opsins in retinas of semi-aquatic snakes belonging to the *Helicops* genus. This convergent diversification of gene copies in independent aquatic snake lineages may suggest a trend in sensory adaptation following secondary aquatic transitions. The degree of spectral complexity observed in aquatic environments is unique; terrestrial ecosystems are not subject to light attenuation and spectral filtering with depth and turbidity, or variable light polarisation by surface refraction. Achieving adequate visual perception in these aquatic environments may therefore prove more challenging than in the static spectral conditions encountered during a dim-light terrestrial ancestry. We suggest that both *Hydrophis* and *Helicops* have convergently expanded their functional opsin repertoires in response to their new and highly variable photic environments. This presents a striking contrast to numerous reports of rapid opsin pseudogenization during transitions to dim-light or aquatic environments in other vertebrates (Fasick et al. 1998; Peichl & Moutairou 1998; Levenson et al. 2006; Wakefield et al. 2008; Jacobs 2009; Gerkema et al. 2013; Jacobs, 2013; Emerling & Springer, 2014; Borges et al. 2015; Simões et al. 2015; Sadier et al. 2018; Gower et al. 2021), including the ancestral snake (Walls 1940 & 1942; Simões et al. 2015; Hsiang et al. 2015). To our knowledge, SWS1 gene duplications have not occurred in terrestrial snakes despite sharing some of the genomic elements that may have facilitated the *Hydrophis* expansions (supplementary figs.S6-11). It is therefore possible that these expansions confer adaptive benefits which are specific to aquatic environments.

However, it is not clear what selection pressures *within* aquatic environments might be associated with SWS1 expansions in these snakes. No particular trait, or combination of traits, distinguish *Hydrophis* and *Helicops* species with multiple SWS1 copies from their single-copy relatives. Species of *Helicops* and the clade spanning *H. cyanocinctus* and *H. spiralis* are primarily nocturnal/crepuscular (Simões et al. 2020), while *H. curtus* is strongly day active (Udyawer et al. 2015); *Helicops* and *H. curtus* strike prey in open water (de Aguiar & Schmidt 2004; Voris & Voris 1983; Udyawer et al. 2015), whereas the other multiple-copy *Hydrophis* are crevice or burrow foragers (Voris & Voris 1983; Rezaie-Atagholipour et al. 2013; Buzas et al. 2019). Dedicated behavioural and observational studies are required to better understand the adaptive significance of SWS1 expansions in aquatic snakes.

## Conclusions

Our study has uncovered the highly dynamic evolution of SWS1 in *Hydrophis* sea snakes, with expansions followed by differential gene losses, or multiple independent gains and fewer losses. Expansions showed close proximity to chromatid ends, and in every instance were followed by functional divergence in wavelength sensitivity at a key spectral tuning site. Retinal transcriptomes revealed simultaneous expression of these spectral subtypes, providing compelling evidence that SWS1 expansions confer novel adaptive phenotypes. Future studies should determine the capacity for increased chromatic distinction in *Hydrophis* using single-cell transcriptomics, RNA fluorescence in situ hybridization, and behavioural studies.

## Materials and Methods

### Taxon Sampling

Specimens and sequences used for whole-genome sequencing, transcriptome analysis and Sanger sequencing were obtained from a wide variety of sources. Localities, sampling and accession numbers are given in supplementary table.S1. Eyes of *H. cyanocinctus* and *Hydrophis atriceps* used for transcriptome analysis were sourced from snakes found in wet markets in both Song Doc town in the Cà Mau province and Vũng Tàu City in the Bà Rịa-Vũng Tàu Province of Vietnam. Eyes of *Hydrophis kingii*, *Hydrophis major*, *Hydrophis peronii*, *Aipysurus laevis*, *Aipysurus mosaicus* and *Hydrelaps darwiniensis* were sourced from snakes caught off coastal Broome, Western Australia. Snakes were euthanized prior to sampling and in accordance with approved animal ethics procedures (University of Adelaide Animal Ethics Committee (S-2015-119). Following euthanasia, whole eyes were immediately dissected using sterile instruments and stored in RNAlater to prevent degradation of RNA. They were then transferred to a -20℃ freezer until extraction. Liver samples from all specimens were collected due to their high DNA yield following extraction and were stored in 100% ethanol in a -20℃ freezer.

### Sequence Data Generation

Nanopore sequence reads and MGI short-read sequences were generated using the same methods detailed in Ludington et al. (2023). Briefly, high molecular weight (HMW) DNA from *Aipysurus laevis* was extracted from 25-50µl of blood using the Circulomics Nanobind CBB Big DNA Kit following the manufacturer’s instructions. Sequence libraries were prepared using the Ultra-Long DNA sequencing kit (SQK-ULK001) and were sequenced across 2 x promethION (FLO-PRO002) flow cells, each for 72 hours. To increase sequencing throughput, DNAse washes (EXP-WSH003) were performed at 24 and 48 hours to unblock pores. Basecalling was then performed using Guppy (v6.1.5) to generate the final sequence data.

Paired end sequence libraries were prepared from tail DNA for 45 samples (supplementary table.S1), with libraries being prepared at both the South Australian Genomic Center (SAGC) and the Ramaciotti Centre for Genomics (RCG). For samples sequenced at the SAGC, DNA was extracted following the Gentra Puregene Tissue Kit protocol, with libraries then being prepared according to the Illumina DNA Prep (M) (Part No. 10000000254 v10) and Illumina to MGI Library Conversion (MGIEasy Universal Library Conversion Kit, Part No. MGI000004155) guidelines, before being sequenced on a MGI DNBSEQ-G400 (2×150bp paired-end) machine. Samples sequenced at RCG had their libraries prepared using the Illumina DNA PCR-Free prep kit before being sequenced on an Illumina NovaSeq 6000 S4 flow cell (2×150bp paired-end). All short-read sequence data were filtered for contaminants and low quality reads against Kraken 2 (v.2.1.2; Wood et al. 2019) and Fastp (v.0.23.2; Chen 2023), respectively.

### Genome Assembly of *Aipysurus laevis*

Prior to assembly, the genome size of *A. laevis* was estimated from the high-accuracy MGI sequencing data using GenomeScope2 (v2.0, Ranallo-Benavidez et al. 2020). Nanopore long-reads were assembled into primary contigs using Flye (v2.9-b1768; Kolmogorov et al. 2019), followed by a single round of Nanopore polishing using Medaka (v1.8.0). Nextpolish (v1.4.1; Hu et al. 2020) was then used to perform two additional rounds of genome polishing using the MGI short-read data. Genome completeness was assessed using BUSCO (v5.2.2; Manni et al. 2021) with the tetrapoda (odb10) database, while genome quality was measured using consensus quality (QV) and k-mer completeness scores generated by Merqury (v1.3; Rhie et al. 2020).

### Distribution and Origins of SWS1 Expansions in Sea Snakes

#### Determining SWS1 Copy Number in Reference Genomes

Long-read assembled genomes for *H. cyanocinctus*, *H. ornatus*, *H. curtus*, *H. elegans*, *H. major* and *A. laevis* were analysed for SWS1 gene duplications (see supplementary table.S1). An annotated *H. curtus* SWS1 reference sequence (Genbank: MT337755; henceforth referred to as ‘the reference SWS1’) was aligned to these six genomes in Geneious Prime (v.2021.1.1) using Minimap2 (Li 2018). Individual exons from the reference SWS1 sequence were also mapped to each genome to check for lone exon duplications, though none were detected. SWS1 copy number, gene orientation and amino acid residues at key spectral tuning sites were then recorded for each species.

#### Estimating SWS1 Copy Number in WGS Samples

SWS1 copy numbers were investigated in 18 species using short-read sequencing: *H. cyanocinctus*, *H. melanocephalus*, *H. atriceps*, *H. spiralis*, *H. darwiniensis*, *H. curtus*, *H. elegans*, *A. laevis*, *H. major*, *H. ocellatus*, *H. ornatus*, *H. peronii*, *H. platurus*, *H. schistosus*, *H. stokesii*, *H. kingii*, *N. naja* and *N. scutatus*. Sequence reads were aligned to the *H. major* reference genome (JAUCBL000000000) using BWA2 (v.2.2.1; Vasimuddin et al. 2019), requiring a minimum mapping quality of 20, before marking duplicates using Sambamba (v0.8.2; Tarasov et al. 2015). *H. major* was chosen as the reference genome because it has a single copy of SWS1 and high assembly accuracy. The program Mosdepth (v0.3.3; Pedersen & Quinlan 2018) was then used to estimate genome-wide coverage statistics for each sample, while SAMtools (v.1.10; Danecek et al. 2021) depth (-a -r chr4:213702160-213714105) was used to estimate the per-base sequence coverage across the SWS1 gene region. For each sample, the SWS1 coverage value was normalised against the average sequencing coverage of chromosome 4. Normalised values were visualised using ggplot2 (v.3.4.2; Wickham 2016) in RStudio (v.2022.12.0; R Core Team 2021) and data was smoothed by loess local regression (span = 1). If SWS1 gene loci had at least double the coverage of the rest of the chromosome, this was interpreted as evidence of duplication under the expectation that reads from multiple copies will align to the single gene locus in *H. major*. To validate the use of normalised coverage in estimating gene copy numbers, we applied our method to the 5,045 ‘complete’ BUSCO genes found in the *H. major* genome assembly. BUSCOs are evolutionarily conserved single-copy orthologs for a given taxonomic group. Consequently, this set of genes acts as a control group for which we expect to see approximately one-fold normalised coverage for each gene in each sample assuming adequate breadth and depth of coverage. For each resequencing sample we calculated their average normalised depth for each of these genes, before visualising their gene coverage distributions using box-plots.

#### SWS1 Exon 1 Sequencing

Genomic DNA was extracted from liver tissue using the Gentra Puregene extraction kit (Qiagen) following the manufacturers’ instructions. DNA products were amplified by touchdown PCR with primers in alignment with the elapid *OPN1SW* exon 1 region (bp136-346; designed by Simões et al. 2020). Primer details are as listed below:

> SWS1F_Elapidae86 -CCATCTTCATGGGCCTTGTCT-

> SWS1R_Elapidae86 -CTGCCACTGTGCCTAGGAAG-

PCR protocol follows that described in Simões et al. (2020). PCR products were run on a 1% agarose gel and sequenced in both directions by an automated sequencer at the Australian Genome Research Facility (AGRF) Adelaide branch. Sequences were assembled in Geneious Prime (v.2021.1.1) by trimming the low-quality ends of each raw sequence. Consensus sequences were created by combining forward and reverse sequences and aligned to the reference SWS1. Indels and low-quality base calls were deleted and chromatogram peaks were checked at spectral tuning site 86 using the heterozygote plugin. Single peaks would indicate the presence of a single SWS1 gene copy (or multiple copies that are homozygous for the site 86 allele). Double peaks of A and T nucleotides may instead indicate gene duplication if these are present in a higher proportion of individuals than expected under Hardy-Weinberg equilibrium (see supplementary fig.S1 for examples). Numbers of expected alleles and Hardy–Weinberg proportions are often used to identify paralogous loci for exclusion from population genetic analyses (e.g. Hohenlohe et al. 2011). Here, double-peak chromatograms at site 86 in more than 60% of at least eight sampled individuals were tentatively inferred to represent the presence of multiple spectrally-distinct SWS1 copies, as this high proportion is observed in *H. cyanocinctus* (see table.S1) for which whole-genome data has confirmed multiple copies (Rossetto et al. 2023).

#### Analysis of Variant Sites within SWS1 Genes

To check for polymorphism at spectral tuning site 86 within single SWS1 gene copies, variants were called across the gene region from the *H. major* alignment files. BCFtools mpileup was used to estimate genotype likelihoods, keeping sites where the mapping and base quality were both greater than 20 (-q 20 -Q 20), before calling variants using the updated multiallelic caller (bcftools call -m; Danecek et al. 2021). Filters were applied to exclude any variants within 10 bases of an insertion or deletion (bcftools filter --SnpGap 10), before filtering for bi-allelic variants with zero missingness across all samples, a minimum coverage of 10 reads and quality 20 using VCFtools (--recode – recode-INFO-all –min-alleles 2 –max-alleles 2 –max-missing 1 –minDP 10 –minQ 20). Amino acid variation at the spectral tuning site was noted (supplementary table.S4).

#### Phylogenetic Origins of the SWS1 Expansion

The evolution of SWS1 duplications and possible losses was reconstructed by mapping inferred copy numbers and spectral tuning at site 86 for 33 sea snake species to the sea snake phylogeny (modified from Sanders et al. 2008 and Ludington et al. 2023 to include all lineages investigated here). Species for which no opsin data was available were removed from the phylogeny. Spectral tuning of each species’ SWS1 opsin was inferred from Simões et al. (2020) and Rossetto et al. (2023) based on MSP data. SWS1 gene copy number was labelled for each species as deduced from works conducted in the current study (figs.1-2). To detect pseudogenes, the SWS1 reference was aligned to reference genomes with Minimap2 (Li 2018) using low-stringency parameters (override k-mer length = 8).

#### Phylogenetic Analysis of SWS1 Sequences

SWS1 gene trees were constructed using sequences extracted from reference genomes for *A. laevis*, *H. ornatus*, *H. curtus*, *H. elegans*, *H. major* and *H. cyanocinctus*. Full sequences (including all exons and introns) were then aligned with the MUSCLE plugin (Edgar 2004) in Geneious Prime (v.2021.1.1) before manual curation. The FASTA alignment was trimmed with ClipKIT (v.2.1.0; Steenwyk et al. 2020) using default parameters. The maximum likelihood SWS1 gene tree was built using the RAxML plugin (v.8.2.12; Stamatakis 2014) with two separate partitions for exons and introns. The *A. laevis* sequence was used as the outgroup because *Aipysurus* and *Hydrophis* are reciprocally monophyletic based on morphological (Rasmussen et al. 2002) and molecular (Sanders et al. 2013) evidence. Selective constraint of the SWS1 genes can be estimated using the ratio of nonsynonymous (dN) over synonymous (dS) substitutions, known as the dN/dS ratio or ω (Nielsen & Yang 1998). However, we decided that this analysis would not be worthwhile, as we believe one of the SWS1 gene copies to be an artefact of mis-assembly (as discussed in Rossetto et al. 2023). Estimating dN/dS ratios may therefore lead us to make false conclusions regarding copy-specific selective pressures.

### Molecular Origin of the SWS1 Expansion

#### Identification of Telomeric Repeat Proximity to SWS1

The proximity of SWS1 genes to telomeric repeat sequences was estimated to explore the possibility that their duplication occurred by unequal sister chromatid exchange during meiosis. The telomere identification toolkit (tidk; https://github.com/tolkit/telomeric-identifier) was used to locate potential telomeric repeats on the chromosome containing SWS1 of genomes for sea snakes *H. cyanocinctus*, *H. curtus*, *H. major*, *H. ornatus*, and on the contig containing the SWS1 gene in the *A. laevis* genome, as well as genomes for terrestrial snakes *Ahaetulla prasina, Bungarus multicinctus, Erythrolamprus reginae, Liasis olivaceus and Vipera latastei* for comparison. SWS1 gene positions were located in these terrestrial species by aligning the reference SWS1 sequence to genomes in Geneious Prime (v.2021.1.1) using Minimap2 (Li 2018). Because it is not known whether a sea snake-specific telomere repeat sequence exists, the standard vertebrate telomere sequence (repeats of TTAGGG) was assumed. This sequence is confirmed to be the telomeric repeat in birds, amphibians, mammals, bony fish and reptiles, including some snakes in the genus *Crotalus* (Meyne et al. 1989). The chromosome or contig containing the SWS1 sequence(s) for each species was used for telomere searching and repeat matches were plotted with tidk.

#### Analysis of Repeat Elements within Snake Genomes

A combined repeat library consisting of *H. elegans* repeat sequences (Ludington et al. 2023) and the curated RepBase repeat library (v2018-10-26; Bao et al. 2015) were used to perform homology based annotation for *H. cyanocinctus*. Repeat information for *H. major* and *H. curtus* was obtained from Ludington et al. (2023). Repeat elements detected in close proximity to duplicated segments encompassing the SWS1 genes of interest were noted.

### Retinal Expression of SWS1 Copies

#### Purification of RNA and Transcriptome Sequencing

Retinal transcriptomes were generated to determine the expression of SWS1 gene subtypes. Two individuals of each *H. atriceps* and *H. cyanocinctus* were selected for eye RNA extraction as these species possess multiple SWS1 gene copies (Rossetto et al. 2023, and this study) and samples were readily obtainable. To compare against species with single SWS1 gene copies, one individual from each of *A. mosaicus*, *A. laevis*, *H. darwiniensis*, *H. kingii*, *H. major* and *H. peronii* were also selected for eye RNA extraction. For each individual, one eye was dissected and the retina homogenized in TRIzol reagent according to the manufacturer’s protocol. Extracted RNA was then purified using the Invitrogen TRIzol Plus RNA Purification Kit (ThermoFisher Scientific) according to the manufacturer’s protocol. RNA product was kept in a -80°C freezer in between practical procedures to avoid degradation. Purified RNA product was sent to the Australian Genomic Research Facility (AGRF) Adelaide branch for quality control using an Agilent ScreenTape Assay and qPCR, and sequencing with the Illumina NovaSeq 6000 system using the SP Reagent Kit.

Trinity (v.2.8.5; Grabherr et al. 2011) was used to *de novo* assemble the Illumina sequence data into transcripts. Prior to assembly, reads were adapter and quality trimmed by Trimmomatic (v0.39; Bolger et al. 2014) using the default parameters specified by Trinity. Following assembly, the reference SWS1 sequence was searched against the assembled transcripts using BLAST (v2.12.0; Altschul et al. 1990) to identify SWS1 transcripts, with candidate SWS1 sequences requiring at least 80% identity and query coverage of the reference SWS1 sequence. Following annotation of SWS1 transcripts, RNA libraries were mapped back to their respective transcriptome assemblies using HISAT2 (v.2.1; Kim et al. 2019) using default parameters, with the resulting alignment files being sorted and indexed using SAMtools (v.1.10; Danecek et al. 2021). Allelic depth of phenylalanine (F) and tyrosine (Y) at site 86 in the SWS1 transcript was then estimated from the aligned data. If either amino acid made up a substantial portion (≥20%) of the total read count at this site, then we excluded the possibility that this was due to sequencing error.

## Data Availability

The data underlying this article are all publicly available. The *Aipysurus laevis* genome assembly has been deposited at GenBank under the accession JBEABB000000000. The version described in this paper is version JBEABB010000000. Sequence data used in the *Aipysurus laevis* genome assembly have been deposited to SRA under the BioProject PRJNA1115802. WGS resequencing and RNA sequencing data have been uploaded to SRA under the accession PRJNA1130540, with per-sample BioSample identifiers detailed in supplementary table.S1. The 104 sanger sequences included in this study have been deposited to GenBank; please see supplementary table.S1 for accession information.

## Acknowledgements

This work was supported by the Australian Research Council Discovery Grant (DP180101688) awarded to KLS and BFS. The lead author IHR was supported by a University of Adelaide Research Scholarship during the time of work.

## References

Altschul SF, Gish W, Miller W, Myers EW, Lipman DJ. 1990. Basic local alignment search tool. Journal of Molecular Biology 215:403–410.

Archer S, Hope A, Partridge JC. 1997. The molecular basis for the green-blue sensitivity shift in the rod visual pigments of the European eel. Proceedings of the Royal Society of London. Series B: Biological Sciences 262:289–295.

Bailey J, Gu Z, Clark R, Knut R, Vallente R, Schwartz S, Adams M, Myers E, Li P, Eichler E. 2002. Recent Segmental Duplications in the Human Genome. Science (New York, N.Y.) 297:1003–1007.

Bao W, Kojima KK, Kohany O. 2015. Repbase Update, a database of repetitive elements in eukaryotic genomes. Mobile DNA 6:11.

Bolger AM, Lohse M, Usadel B. 2014. Trimmomatic: a flexible trimmer for Illumina sequence data. Bioinformatics 30:2114–2120.

Borges R, Khan I, Johnson WE, Gilbert MTP, Zhang G, Jarvis ED, O’Brien SJ, Antunes A. 2015. Gene loss, adaptive evolution and the co-evolution of plumage coloration genes with opsins in birds. BMC Genomics 16:751.

Britt LL, Loew ER, McFarland WN. 2001. Visual pigments in the early life stages of Pacific northwest marine fishes. Journal of Experimental Biology 204:2581–2587.

Buzas B, Farkas B, Gulyas E, Geczy C. 2019. The sea snakes (Elapidae: Hydrophiinae) of Fujairah. 26.

Carleton KL, Spady TC, Streelman JT, Kidd MR, McFarland WN, Loew ER. 2008. Visual sensitivities tuned by heterochronic shifts in opsin gene expression. BMC Biology 6:22.

Carlton J, Angiuoli S, Suh B, Kooij T, Pertea M, Silva J, Ermolaeva M, Allen J, Selengut J, Koo H, et al. 2002. Genome sequence and comparative analysis of the model rodent malaria parasite *Plasmodium yoelii yoelii*. Nature 419:512–519.

Chen J-M, Chuzhanova N, Stenson PD, Férec C, Cooper DN. 2005. Intrachromosomal serial replication slippage in trans gives rise to diverse genomic rearrangements involving inversions. Human Mutation 26:362–373.

Chen L, Zhou W, Zhang L, Zhang F. 2014. Genome Architecture and Its Roles in Human Copy Number Variation. Genomics & Informatics 12:136.

Chen S. 2023. Ultrafast one-pass FASTQ data preprocessing, quality control, and deduplication using fastp. iMeta 2:e107.

Corbo J. 2021. Vitamin A1/A2 chromophore exchange: Its role in spectral tuning and visual plasticity. Developmental Biology 475.

Cronin TW, Johnsen S, Marshall NJ, Warrant EJ. 2014. Visual Ecology. In: Princeton University Press.

Dalton BE, Loew ER, Cronin TW, Carleton KL. 2014. Spectral tuning by opsin coexpression in retinal regions that view different parts of the visual field. Proceedings of the Royal Society B: Biological Sciences 281:20141980.

Danecek P, Bonfield JK, Liddle J, Marshall J, Ohan V, Pollard MO, Whitwham A, Keane T, McCarthy SA, Davies RM, et al. 2021. Twelve years of SAMtools and BCFtools. GigaScience 10:giab008.

de Aguiar LFS, Di-Bernardo M. 2004. Diet and Feeding Behavior of *Helicops infrataeniatus* (Serpentes: Colubridae: Xenodontinae) in Southern Brazil. Studies on Neotropical Fauna and Environment 39:7–14.

Dulai KS, von Dornum M, Mollon JD, Hunt DM. 1999. The Evolution of Trichromatic Color Vision by Opsin Gene Duplication in New World and Old World Primates. Genome Research 9:629–638.

Edgar RC. 2004. MUSCLE: multiple sequence alignment with high accuracy and high throughput. Nucleic Acids Research 32:1792–1797.

Eichler E, Sankoff D. 2003. Structural Dynamics of Eukaryotic Chromosome Evolution. Science (New York, N.Y.) 301:793–797.

Emerling CA, Springer MS. 2014. Eyes underground: Regression of visual protein networks in subterranean mammals. Molecular Phylogenetics and Evolution 78:260–270.

Fasick JI, Cronin TW, Hunt DM, Robinson PR. 1998. The visual pigments of the bottlenose dolphin (*Tursiops truncatus*). Visual Neuroscience 15:643–651.

Fujiyabu C, Sato K, Utari NML, Ohuchi H, Shichida Y, Yamashita T. 2019. Evolutionary history of teleost intron-containing and intron-less rhodopsin genes. Scientific Reports 9:10653.

Gerkema MP, Davies WIL, Foster RG, Menaker M, Hut RA. 2013. The nocturnal bottleneck and the evolution of activity patterns in mammals. Proceedings of the Royal Society B: Biological Sciences 280:20130508.

Gower DJ, Fleming JF, Pisani D, Vonk FJ, Kerkkamp HMI, Peichl L, Meimann S, Casewell NR, Henkel CV, Richardson MK, et al. 2021. Eye-Transcriptome and Genome-Wide Sequencing for Scolecophidia: Implications for Inferring the Visual System of the Ancestral Snake. Genome Biology and Evolution 13:evab253.

Grabherr MG, Haas BJ, Yassour M, Levin JZ, Thompson DA, Amit I, Adiconis X, Fan L, Raychowdhury R, Zeng Q, et al. 2011. Full-length transcriptome assembly from RNA-Seq data without a reference genome. Nature biotechnology 29:644–652.

Hagen JFD, Roberts NS, Johnston RJ. 2023. The evolutionary history and spectral tuning of vertebrate visual opsins. Developmental Biology 493:40–66.

Hart NS, Coimbra JP, Collin SP, Westhoff G. 2012. Photoreceptor types, visual pigments, and topographic specializations in the retinas of hydrophiid sea snakes. Journal of Comparative Neurology 520:1246–1261.

Hauzman E, Pierotti MER, Bhattacharyya N, Tashiro JH, Yovanovich CAM, Campos PF, Ventura DF, Chang BSW. 2021. Simultaneous Expression of UV and Violet SWS1 Opsins Expands the Visual Palette in a Group of Freshwater Snakes. Molecular Biology and Evolution 38:5225–5240.

Hohenlohe PA, Amish SJ, Catchen JM, Allendorf FW, Luikart G. 2011. Next-generation RAD sequencing identifies thousands of SNPs for assessing hybridization between rainbow and westslope cutthroat trout. Molecular Ecology Resources 11:117–122.

Horvath J, Bailey J, Locke D, Eichler E. 2001. Lessons from the human genome: Transitions between euchromatin and heterochromatin. Human Molecular Genetics 10.

Hsiang AY, Field DJ, Webster TH, Behlke ADB, Davis MB, Racicot RA, Gauthier JA. 2015. The origin of snakes: revealing the ecology, behavior, and evolutionary history of early snakes using genomics, phenomics, and the fossil record. BMC Evolutionary Biology 15:87.

Hu J, Fan J, Sun Z, Liu S. 2020. NextPolish: a fast and efficient genome polishing tool for long-read assembly. Bioinformatics 36:2253–2255.

Hunt DM, Dulai KS, Cowing JA, Julliot C, Mollon JD, Bowmaker JK, Li W-H, Hewett-Emmett D. 1998. Molecular evolution of trichromacy in primates. Vision Research 38:3299–3306.

Jacobs GH. 2009. Evolution of colour vision in mammals. Philosophical Transactions of the Royal Society B: Biological Sciences 364:2957–2967.

Jacobs GH. 2013. Losses of functional opsin genes, short-wavelength cone photopigments, and color vision—A significant trend in the evolution of mammalian vision. Visual Neuroscience 30:39–53.

Kim D, Paggi JM, Park C, Bennett C, Salzberg SL. 2019. Graph-based genome alignment and genotyping with HISAT2 and HISAT-genotype. Nature Biotechnology 37:907–915.

Kolmogorov M, Yuan J, Lin Y, Pevzner PA. 2019. Assembly of long, error-prone reads using repeat graphs. Nature Biotechnology 37:540–546.

Lagman D, Ocampo Daza D, Widmark J, Abalo XM, Sundström G, Larhammar D. 2013. The vertebrate ancestral repertoire of visual opsins, transducin alpha subunits and oxytocin/vasopressin receptors was established by duplication of their shared genomic region in the two rounds of early vertebrate genome duplications. BMC Evolutionary Biology 13:238.

Larhammar D, Nordström K, Larsson TA. 2009. Evolution of vertebrate rod and cone phototransduction genes. Philosophical Transactions of the Royal Society B: Biological Sciences 364:2867–2880.

Levenson DH, Ponganis PJ, Crognale MA, Deegan JF, Dizon A, Jacobs GH. 2006. Visual pigments of marine carnivores: pinnipeds, polar bear, and sea otter. Journal of Comparative Physiology A 192:833–843.

Li A, Wang J, Sun K, Wang S, Zhao X, Wang T, Xiong L, Xu W, Qiu L, Shang Y, et al. 2021. Two Reference-Quality Sea Snake Genomes Reveal Their Divergent Evolution of Adaptive Traits and Venom Systems. Molecular Biology and Evolution 38:4867–4883.

Li H. 2018. Minimap2: pairwise alignment for nucleotide sequences. Bioinformatics 34:3094–3100.

Lin J-J, Wang F-Y, Li W-H, Wang T-Y. 2017. The rises and falls of opsin genes in 59 ray-finned fish genomes and their implications for environmental adaptation. Scientific Reports 7:15568.

Ludington AJ, Hammond JM, Breen J, Deveson IW, Sanders KL. 2023. New chromosome-scale genomes provide insights into marine adaptations of sea snakes (Hydrophis: Elapidae). BMC Biol.

Luehrmann M, Stieb SM, Carleton KL, Pietzker A, Cheney KL, Marshall NJ. 2018. Short-term colour vision plasticity on the reef: changes in opsin expression under varying light conditions differ between ecologically distinct fish species. Journal of Experimental Biology 221:jeb175281.

Mable BK. 2004. ‘Why polyploidy is rarer in animals than in plants’: myths and mechanisms. Biological Journal of the Linnean Society 82:453–466.

Manni M, Berkeley MR, Seppey M, Simão FA, Zdobnov EM. 2021. BUSCO Update: Novel and Streamlined Workflows along with Broader and Deeper Phylogenetic Coverage for Scoring of Eukaryotic, Prokaryotic, and Viral Genomes. Molecular Biology and Evolution 38:4647–4654.

Mefford H, Trask B. 2002. Mefford HC, Trask BJ. The complex structure and dynamic evolution of human subtelomeres. Nat Rev Genet 3: 91–102. Nature reviews. Genetics 3:91-102.

Meyer A, Van de Peer Y. 2005. From 2R to 3R: evidence for a fish-specific genome duplication (FSGD). BioEssays 27:937–945.

Meyne J, Ratliff RL, Moyzis RK. 1989. Conservation of the Human Telomere Sequence (TTAGGG)n among Vertebrates. Proceedings of the National Academy of Sciences of the United States of America 86:7049–7053.

Musilova Z, Cortesi F, Matschiner M, Davies WIL, Patel JS, Stieb SM, de Busserolles F, Malmstrøm M, Tørresen OK, Brown CJ, et al. 2019. Vision using multiple distinct rod opsins in deep-sea fishes. Science 364:588–592.

Musilova Z, Salzburger W, Cortesi F. 2021. The Visual Opsin Gene Repertoires of Teleost Fishes: Evolution, Ecology, and Function. Annual Review of Cell and Developmental Biology 37:441–468.

Nathans J, Thomas D, Hogness DS. 1986. Molecular Genetics of Human Color Vision: The Genes Encoding Blue, Green, and Red Pigments. Science 232:193–202.

Nielsen R, Yang Z. 1998. Likelihood Models for Detecting Positively Selected Amino Acid Sites and Applications to the HIV-1 Envelope Gene. Genetics 148:929–936.

Nordström K, Larsson TA, Larhammar D. 2004. Extensive duplications of phototransduction genes in early vertebrate evolution correlate with block (chromosome) duplications. Genomics 83:852–872.

Otto SP, Whitton J. 2000. Polyploid incidence and evolution. Annual review of genetics 34:401–437.

Pan W, Byrne-Steele M, Wang C, Lu S, Clemmons S, Zahorchak RJ, Han J. 2014. DNA polymerase preference determines PCR priming efficiency. BMC Biotechnology 14:10.

Passananti C, Davies B, Ford M, Fried M. 1987. Structure of an inverted duplication formed as a first step in a gene amplification event: implications for a model of gene amplification. The EMBO Journal 6:1697–1703.

Pedersen BS, Quinlan AR. 2018. Mosdepth: quick coverage calculation for genomes and exomes. Bioinformatics 34:867–868.

Pegueroles C, Laurie S, Albà MM. 2013. Accelerated Evolution after Gene Duplication: A Time-Dependent Process Affecting Just One Copy. Molecular Biology and Evolution 30:1830–1842.

Peichl L, Moutairou K. 1998. Absence of short-wavelength sensitive cones in the retinae of seals (Carnivora) and African giant rats (Rodentia). European Journal of Neuroscience 10:2586–2594.

Ranallo-Benavidez TR, Jaron KS, Schatz MC. 2020. GenomeScope 2.0 and Smudgeplot for reference-free profiling of polyploid genomes. Nature Communications 11:1432.

Rasmussen A. 2002. Phylogenetic analysis of the “true” aquatic elapid snakes Hydrophiinae (sensu Smith et al., 1977) indicates two independent radiations into water. Steenstrupia 27:47–63.

Reams AB, Roth JR. 2015. Mechanisms of Gene Duplication and Amplification. Cold Spring Harbor Perspectives in Biology 7.

Rennison DJ, Owens GL, Taylor JS. 2012. Opsin gene duplication and divergence in ray-finned fish. Molecular Phylogenetics and Evolution 62:986–1008.

Rezaie-Atagholipour M, Riyahi-Bakhtiari A, Sajjadi M. 2013. Feeding Habits of the Annulated Sea Snake, *Hydrophis cyanocinctus*, in the Persian Gulf. Journal of Herpetology 47:328–330.

Rhie A, Walenz BP, Koren S, Phillippy AM. 2020. Merqury: reference-free quality, completeness, and phasing assessment for genome assemblies. Genome Biology 21:245.

Rossetto IH, Sanders KL, Simões BF, Van Cao N, Ludington AJ. 2023. Functional Duplication of the Short-Wavelength-Sensitive Opsin in Sea Snakes: Evidence for Reexpanded Color Sensitivity Following Ancestral Regression. Genome Biology and Evolution 15:evad107.

Sadier A, Davies KTJ, Yohe LR, Yun K, Donat P, Hedrick BP, Dumont ER, Dávalos LM, Rossiter SJ, Sears KE. 2018. Multifactorial processes underlie parallel opsin loss in neotropical bats. eLife 7:e37412.

Sanders KL, Lee MSY, Leys R, Foster R, Keogh SJ. 2008. Molecular phylogeny and divergence dates for Australasian elapids and sea snakes (hydrophiinae): evidence from seven genes for rapid evolutionary radiations. Journal of Evolutionary Biology 21:682–695.

Sanders KL, Rasmussen AR, Mumpuni, Elmberg J, de Silva A, Guinea ML, Lee MSY. 2013. Recent rapid speciation and ecomorph divergence in Indo-Australian sea snakes. Molecular Ecology 22:2742–2759.

See DR, Brooks S, Nelson JC, Brown-Guedira G, Friebe B, Gill BS. 2006a. Gene evolution at the ends of wheat chromosomes. Proceedings of the National Academy of Sciences 103:4162–4167.

See DR, Brooks S, Nelson JC, Brown-Guedira G, Friebe B, Gill BS. 2006b. Gene evolution at the ends of wheat chromosomes. Proceedings of the National Academy of Sciences 103:4162–4167.

Seehausen O, Terai Y, Magalhaes I, Carleton K, Mrosso H, Miyagi R, Sluijs I, Schneider M, Maan M, Tachida H, et al. 2008. Speciation through sensory drive in cichlid fish. Nature 455:620–626.

Shay JW, Wright WE. 2019. Telomeres and telomerase: three decades of progress. Nature Reviews Genetics 20:299–309.

Sherratt E, Rasmussen AR, Sanders KL. 2018. Trophic specialization drives morphological evolution in sea snakes. Royal Society Open Science 5:172141.

Simoes B, Sampaio F, Jared C, Antoniazzi M, Loew E, Bowmaker J, Rodriguez A, Hart N, Hunt D, Partridge J, et al. 2015. Visual system evolution and the nature of the ancestral snake. Journal of Evolutionary Biology. 28:12.

Simões BF, Gower DJ, Rasmussen AR, Sarker MAR, Fry GC, Casewell NR, Harrison RA, Hart NS, Partridge JC, Hunt DM, et al. 2020. Spectral Diversification and Trans-Species Allelic Polymorphism during the Land-to-Sea Transition in Snakes. Current Biology.

Simões BF, Sampaio FL, Douglas RH, Kodandaramaiah U, Casewell NR, Harrison RA, Hart NS, Partridge JC, Hunt DM, Gower DJ. 2016. Visual Pigments, Ocular Filters and the Evolution of Snake Vision. Molecular Biology and Evolution 33:2483–2495.

Smallwood PM, Wang Y, Nathans J. 2002. Role of a locus control region in the mutually exclusive expression of human red and green cone pigment genes. Proceedings of the National Academy of Sciences 99:1008–1011.

Stamatakis A. 2014. RAxML version 8: a tool for phylogenetic analysis and post-analysis of large phylogenies. Bioinformatics 30:1312–1313.

Steenwyk JL, Buida TJ, III, Li Y, Shen X-X, Rokas A. 2020. ClipKIT: A multiple sequence alignment trimming software for accurate phylogenomic inference. PLOS Biology 18:e3001007.

Stieb SM, Carleton KL, Cortesi F, Marshall NJ, Salzburger W. 2016. Depth-dependent plasticity in opsin gene expression varies between damselfish (Pomacentridae) species. Molecular Ecology 25:3645–3661.

Tarasov A, Vilella AJ, Cuppen E, Nijman IJ, Prins P. 2015. Sambamba: fast processing of NGS alignment formats. Bioinformatics 31:2032–2034.

Udyawer V, Simpfendorfer CA, Heupel MR. 2015. Diel patterns in three-dimensional use of space by sea snakes. Animal Biotelemetry 3:29.

Vasimuddin M, Misra S, Li H, Aluru S editors. 2019 IEEE International Parallel and Distributed Processing Symposium (IPDPS). 2019 20-24 May 2019.

Voris HK, Voris HH. 1983. Feeding Strategies in Marine Snakes: An Analysis of Evolutionary, Morphological, Behavioral and Ecological Relationships1. American Zoologist 23:411–425.

Wakefield MJ, Anderson M, Chang E, Wei K-J, Kaul R, Graves JAM, GrÜTzner F, Deeb SS. 2008. Cone visual pigments of monotremes: Filling the phylogenetic gap. Visual Neuroscience 25:257–264.

Walls GL. 1940. Ophthalmological Implications for the Early History of the Snakes. Copeia 1940:1–8.

Walls GL. 1942. The vertebrate eye and its adaptive radiation. Cranbrook Institute of Science. In: Cranbrook Press.[aHB, IG].

Ward MN, Churcher AM, Dick KJ, Laver CRJ, Owens GL, Polack MD, Ward PR, Breden F, Taylor JS. 2008. The molecular basis of color vision in colorful fish: Four Long Wave-Sensitive (LWS) opsins in guppies (*Poecilia reticulata*) are defined by amino acid substitutions at key functional sites. BMC Evolutionary Biology 8:210.

Wickham H. 2016. ggplot2: Elegant Graphics for Data Analysis. In: Springer-Verlag New York.

Wood DE, Lu J, Langmead B. 2019. Improved metagenomic analysis with Kraken 2. Genome Biology 20:257.

Yokoyama S. 2002. Molecular evolution of color vision in vertebrates. Gene 300:69–78.

Yokoyama S, Yokoyama R. 2003. Adaptive evolution of photoreceptors and visual pigments in vertebrates. Annual Review of Ecology and Systematics 27:543–567.

