## Supplementary Figures for "Dynamic Expansions and Retinal Expression of Spectrally Distinct Short-Wavelength Opsin Genes in Sea Snakes"

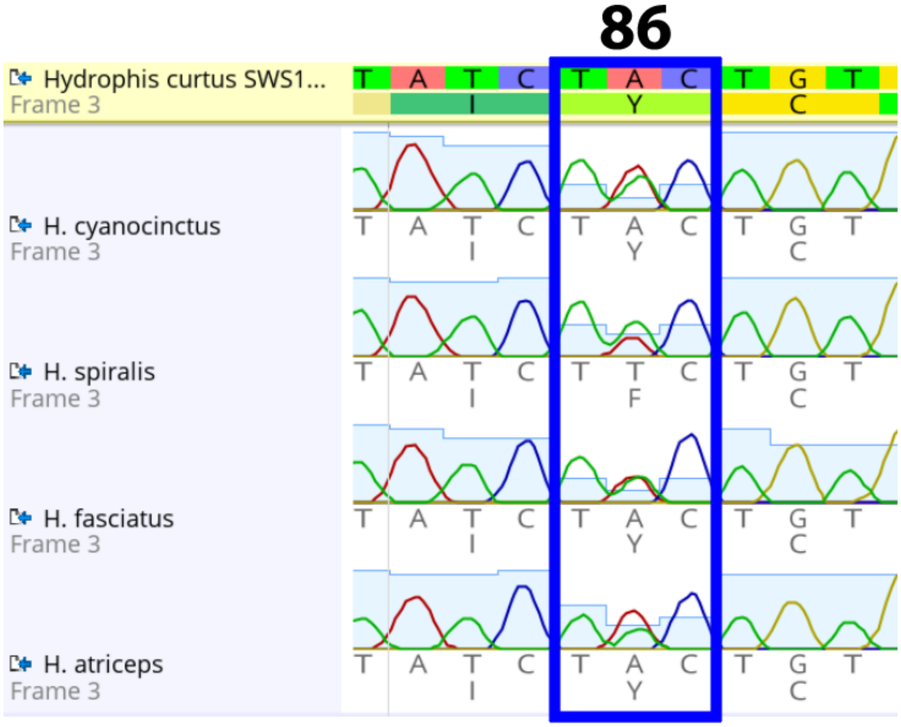


***Figure.S1: Chromatograms at spectral tuning Site 86 of the SWS1 gene in four sea snake species.***
*Green = thymine, Red = adenine, Yellow = guanine, Blue = cytosine. The blue rectangle encompasses amino acid spectral tuning site 86. Chromatograms were generated in Geneious Prime (version 2021.1.1).*

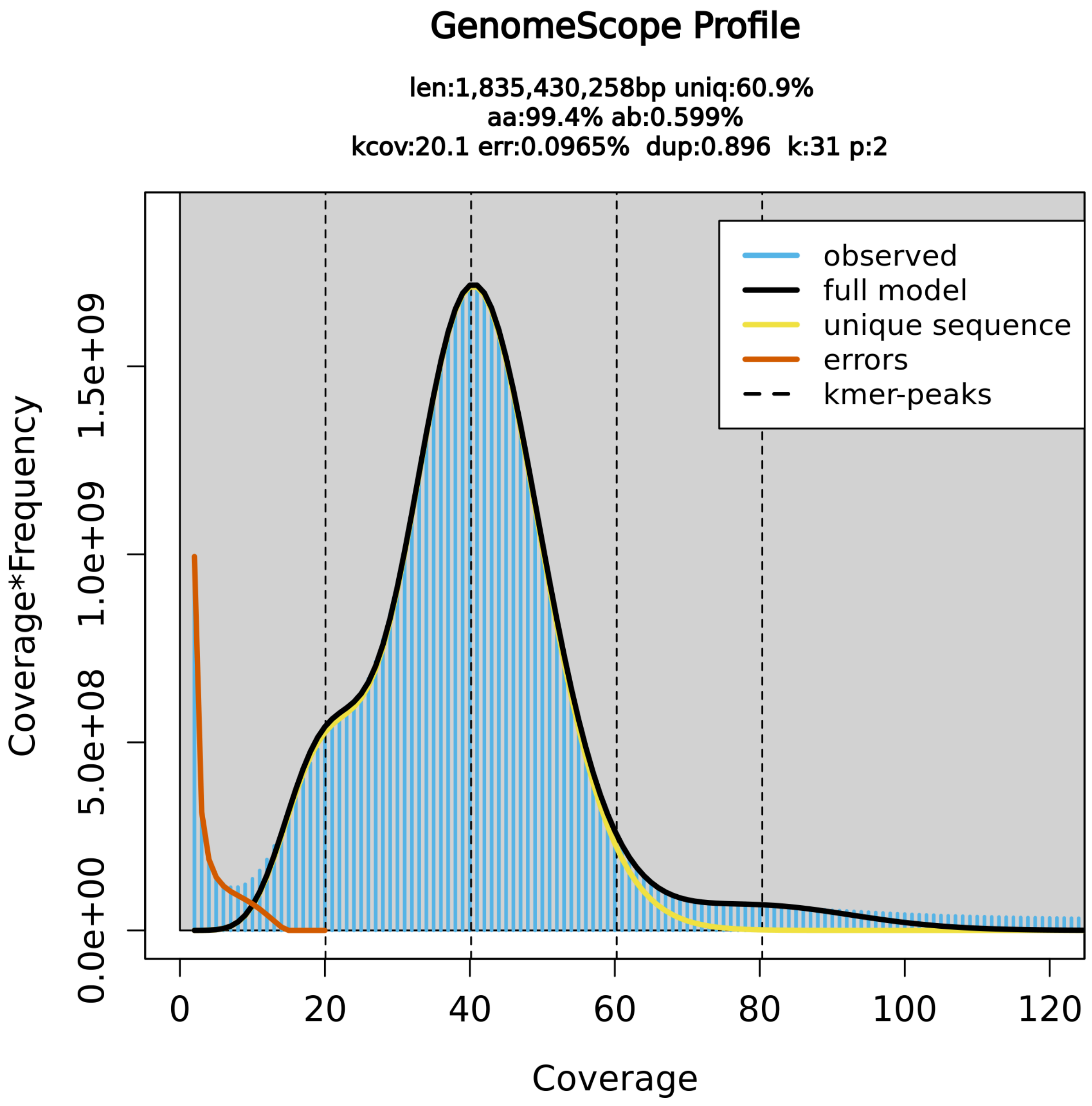


***Figure.S2: Estimated genome size for Aipysurus laevis.*** *K-mer spectra and fitted models generated by GenomeScope2 for A. laevis from short read data using 31-mers. The main central peak represents the homozygous portion of the genome, with the smaller left-shoulder representing the heterozygous sequence. Erroneous K-mers are represented by the peak of low coverage K-mers modelled by the red line.*

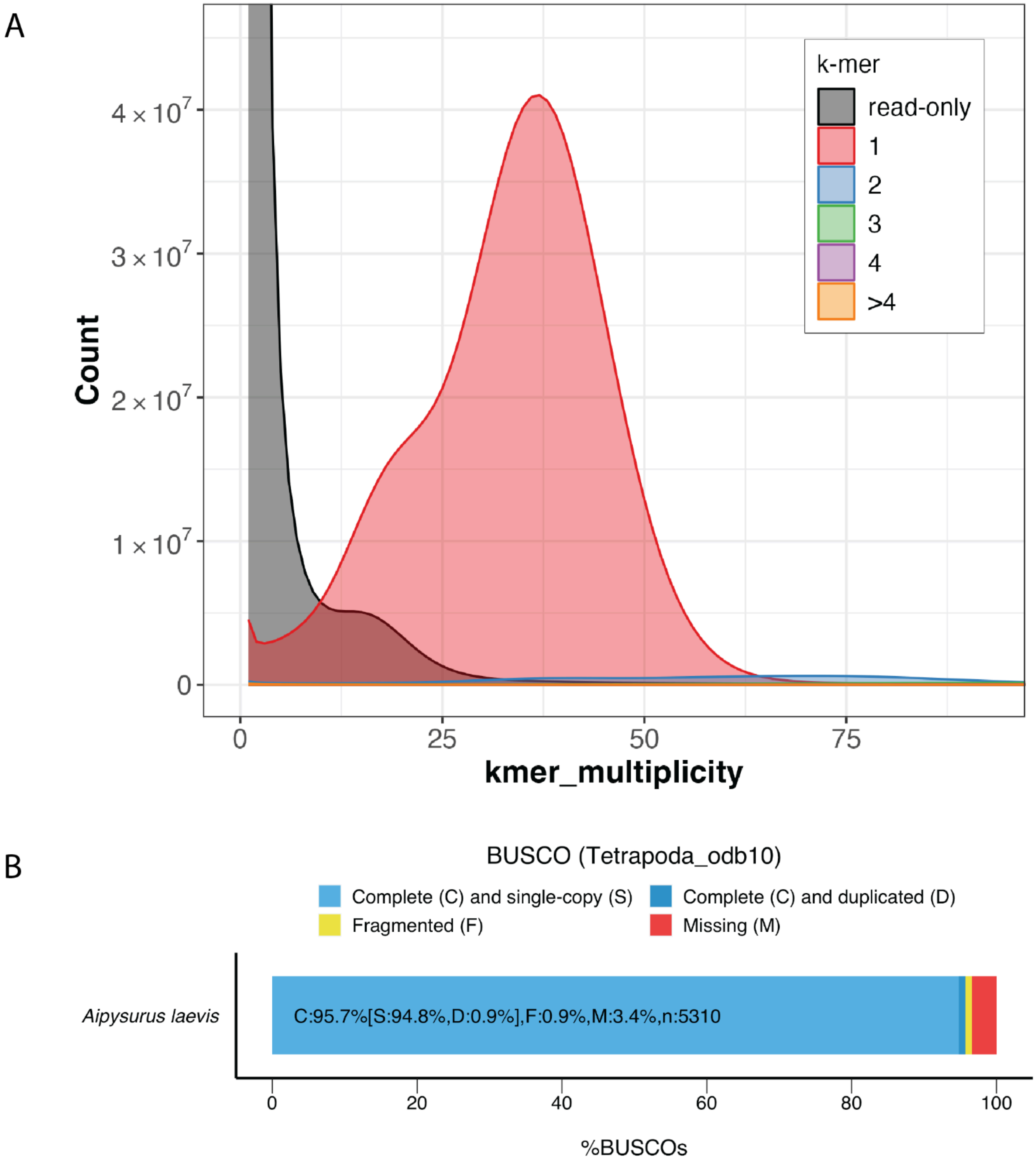


***Figure.S3: i) Aipysurus laevis assembly k-mer spectra and BUSCO completeness results***.

*A) K-mer spectra results generated by Merqury. The x-axis represents k-mer multiplicity, where k = 21, while the y-axis displays the number of k-mers. The black distribution represents k-mers only found in the read-set, with the remaining colours representing how many times read k-mers are found in the assembly, with red representing one time, blue two times, green three times, purple four times and orange more than four times. The shoulder to the left of the red peak represents the heterozygous portion of the genome, with the corresponding shoulder in the black curve representing the alternate sequence that was not included in the haploid genome. The high-frequency, low-multiplicity K-mers in black represent erroneous K-mers. B) Bar plot visualising the BUSCO results using the Tetrapoda odb10 database.*

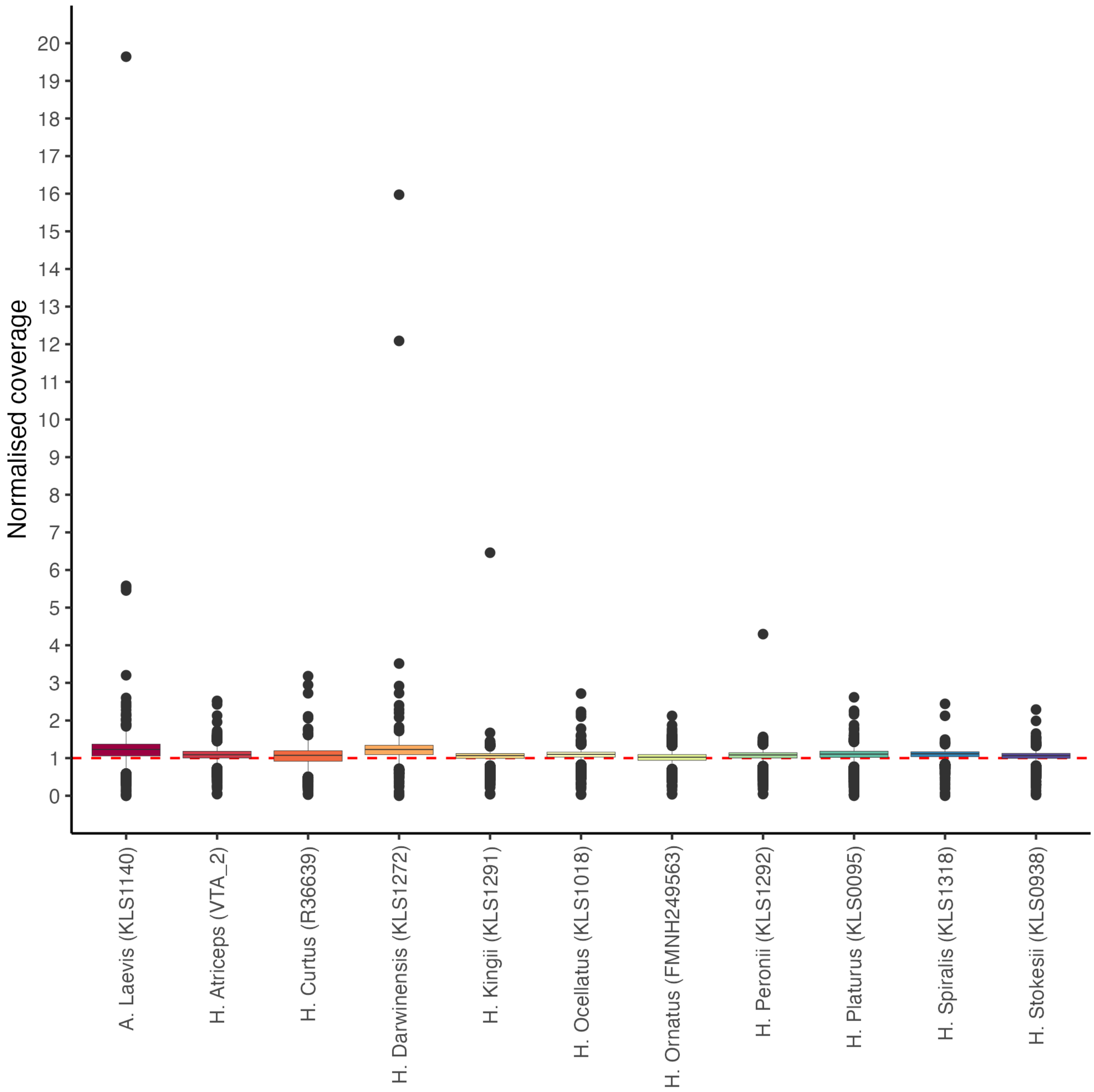


***Figure.S4: Normalised coverage plot of 5,045 BUSCO genes in chromosome four of 11 short-read sea snake genomes.*** *Average normalised coverage values were estimated for each of H. majors 5,045 complete single-copy orthologs for each snake. Across all snakes, the median normalised coverage is close to the expected 1x value, represented by the dashed red line. Within each box-plot, the black horizontal line represents the median value, the upper and lower bounds of the box are the 75th and 25th percentiles, while whiskers extend 1.5 * IQR from the upper and lower bounds. Outliers are shown by the black points extending beyond the whiskers.*

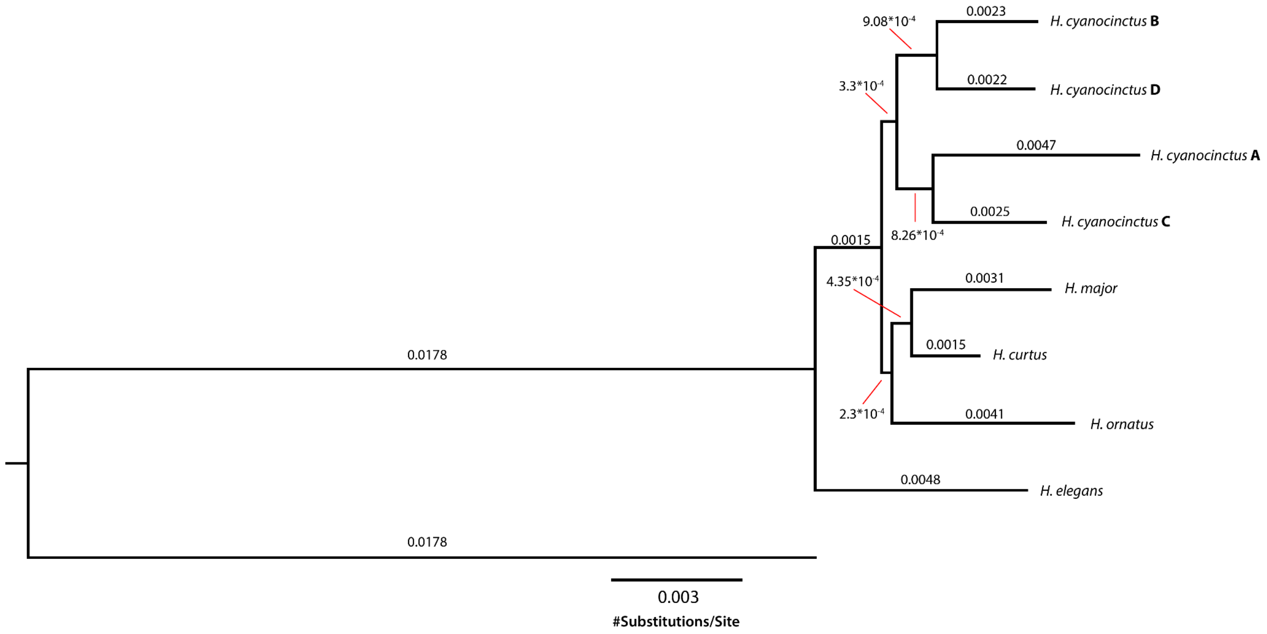


***Figure.S5: Maximum likelihood tree of SWS1 genes from snakes with long-read genomes.*** *Branch labels and scale bar display genetic distance (number of nucleotide substitutions per site). Tree was generated using the RAxML plugin (Stamatakis 2014) in Geneious Prime (v.2021.1.1). The A. laevis sequence was the outgroup. Number of bootstraps generated = 100, parsimony random seed = 1, nucleotide model = GTR-35 GAMMA.*

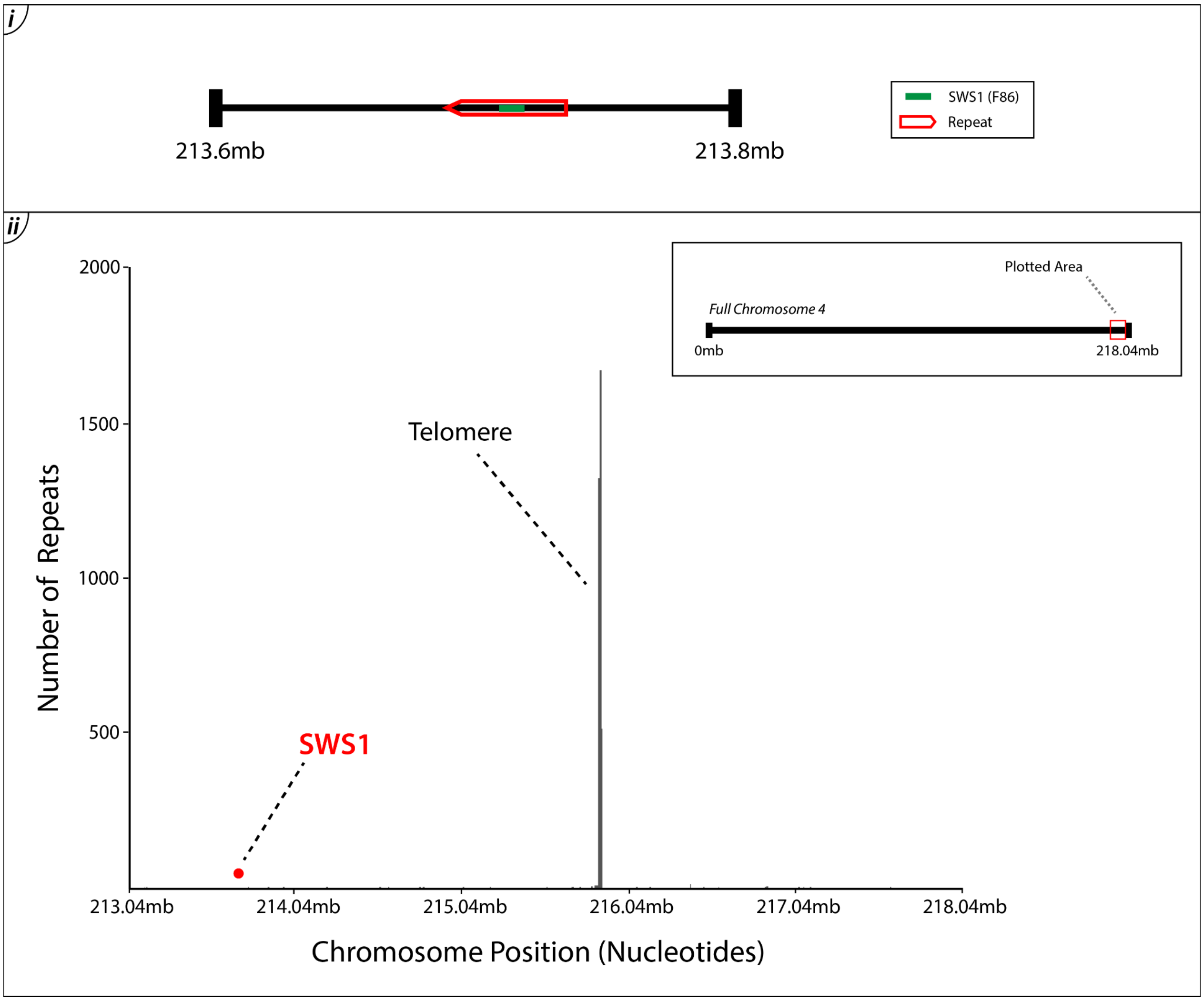


***Figure.S6: i) Location of SWS1 gene in H. major. ii) Telomere proximity to SWS1 gene on chromosome 4 of the H. major genome.*** *SWS1 gene location in the H. major genome (Ludington et al. 2023; Accession = JAUCBL000000000) was inferred from table.S3. Marked repeat areas are homologous with those annotated in H. cyanocinctus (fig.4). The y-axis displays the number of TTAGGG telomere repeat sequences present at a particular chromosome position on the x-axis.*

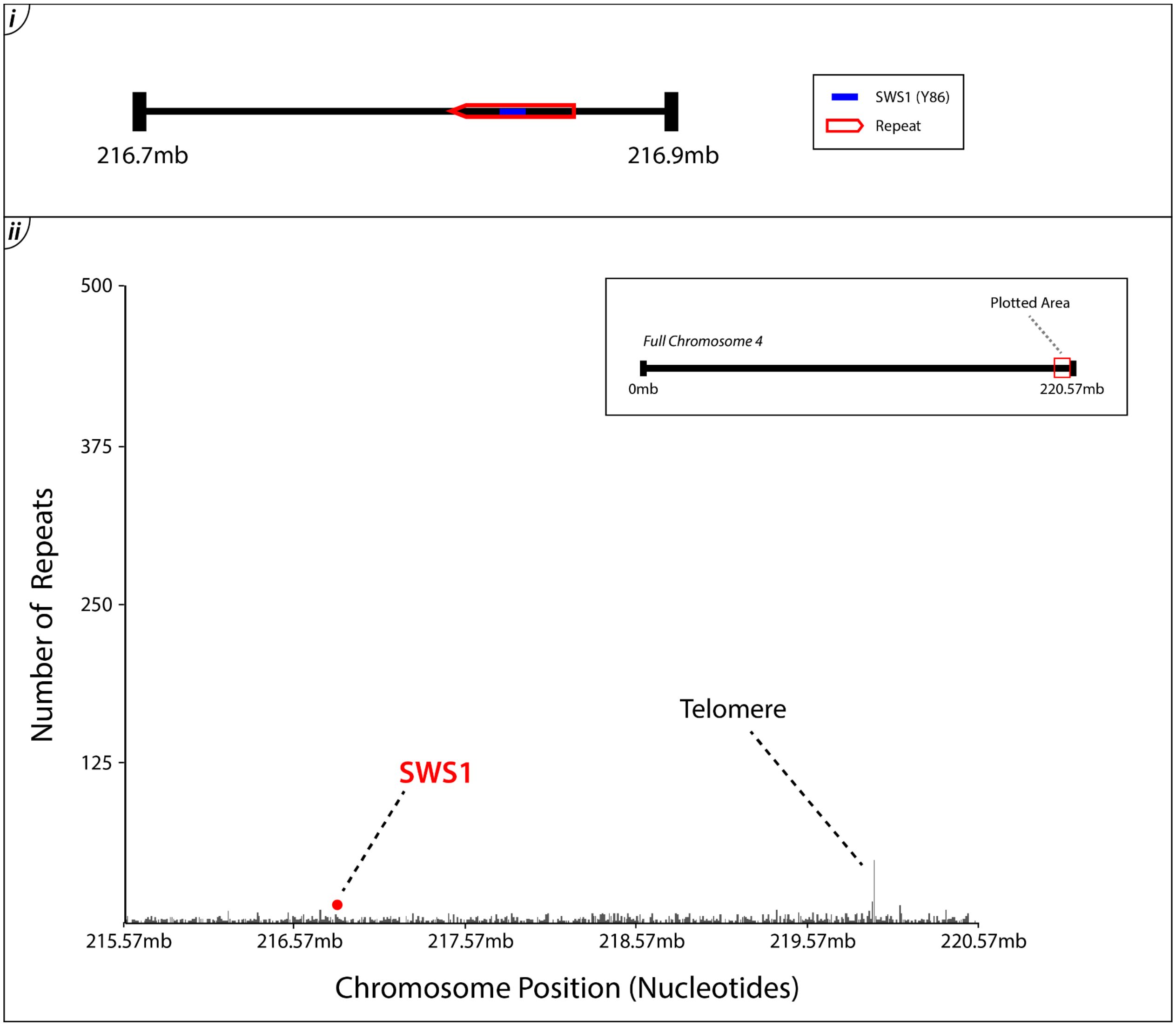


***Figure.S7: i) Location of SWS1 gene in H. curtus. ii) Telomere proximity to SWS1 gene on chromosome 4 of the H. curtus genome.*** *SWS1 gene location in the H. curtus genome (combined data from Li et al. 2021; Genbank = GCA_019473425.1 and Ludington et al. 2023; Accession = SAMN23222007) was inferred from table.S3. Marked repeat areas are homologous with those annotated in H. cyanocinctus (fig.4). The y-axis displays the number of TTAGGG telomere repeat sequences present at a particular chromosome position on the x-axis.*

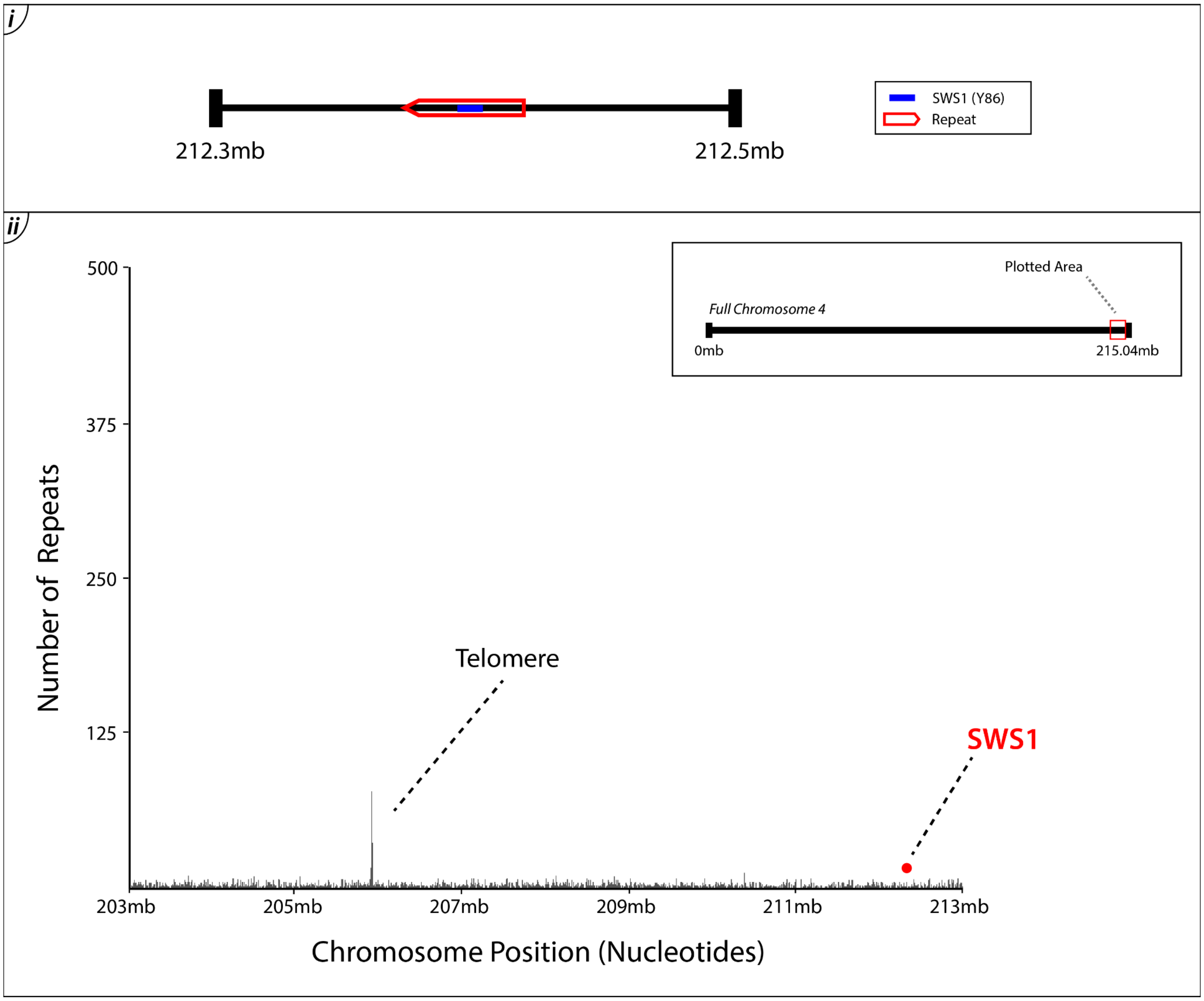


***Figure.S8: i) Location of SWS1 gene in H. ornatus. ii) Telomere proximity to SWS1 gene on chromosome 4 of the H. ornatus genome.*** *SWS1 gene location in the H. ornatus genome (Ludington et al. 2023; Accession= SAMN23222008) was inferred from table.S3. Marked repeat areas are homologous with those annotated in H. cyanocinctus (fig.4). The y-axis displays the number of TTAGGG telomere repeat sequences present at a particular chromosome position on the x-axis.*

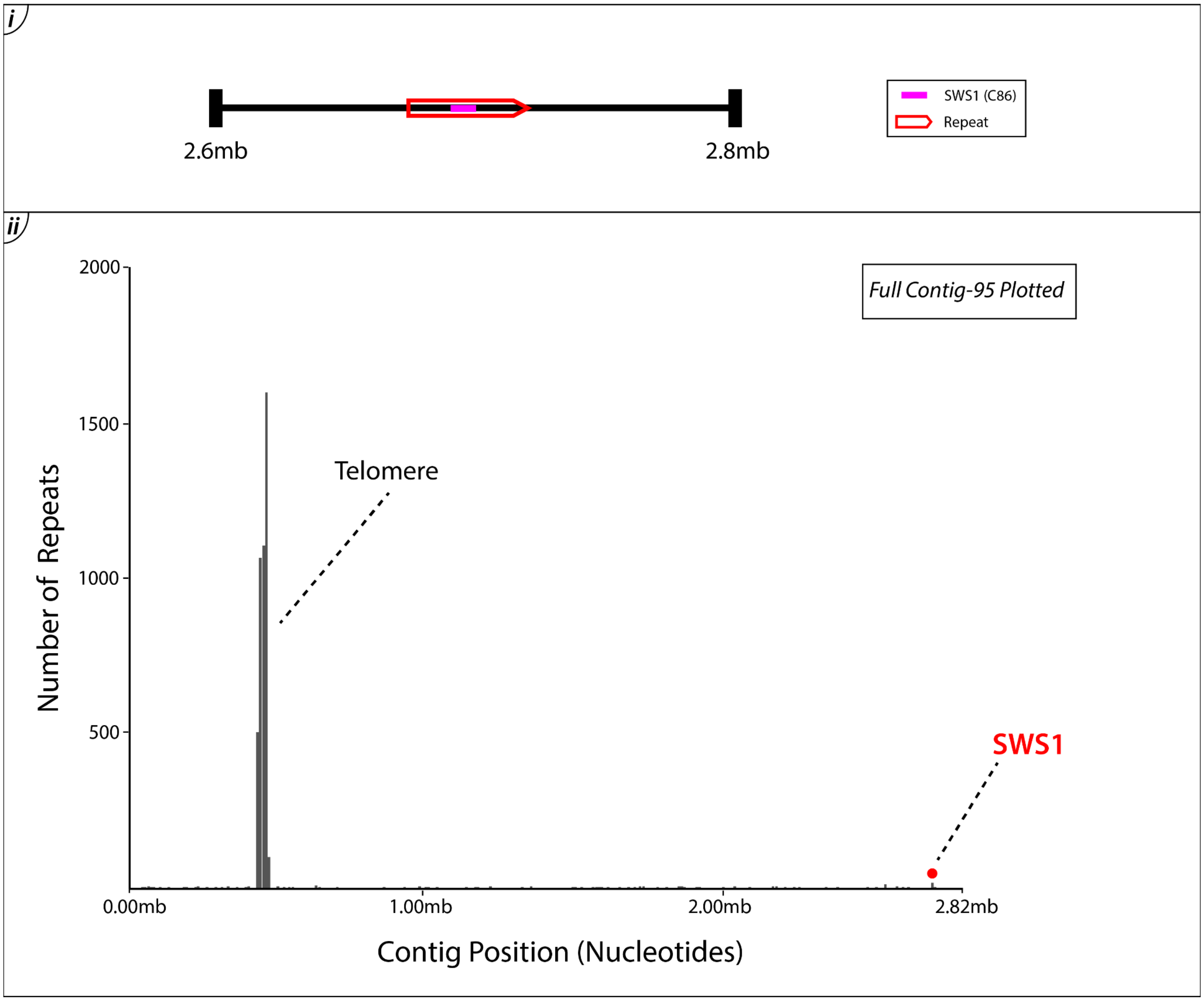


***Figure.S9: i) Location of SWS1 gene in A. laevis. ii) Telomere proximity to SWS1 gene on contig-95 of the A. laevis genome.*** *SWS1 gene location in the A. laevis genome (Accession = PRJNA1115802) was inferred from table.S3. Marked repeat areas are homologous with those annotated in H. cyanocinctus (fig.4). The y-axis displays the number of TTAGGG telomere repeat sequences present at a particular contig position on the x-axis.*

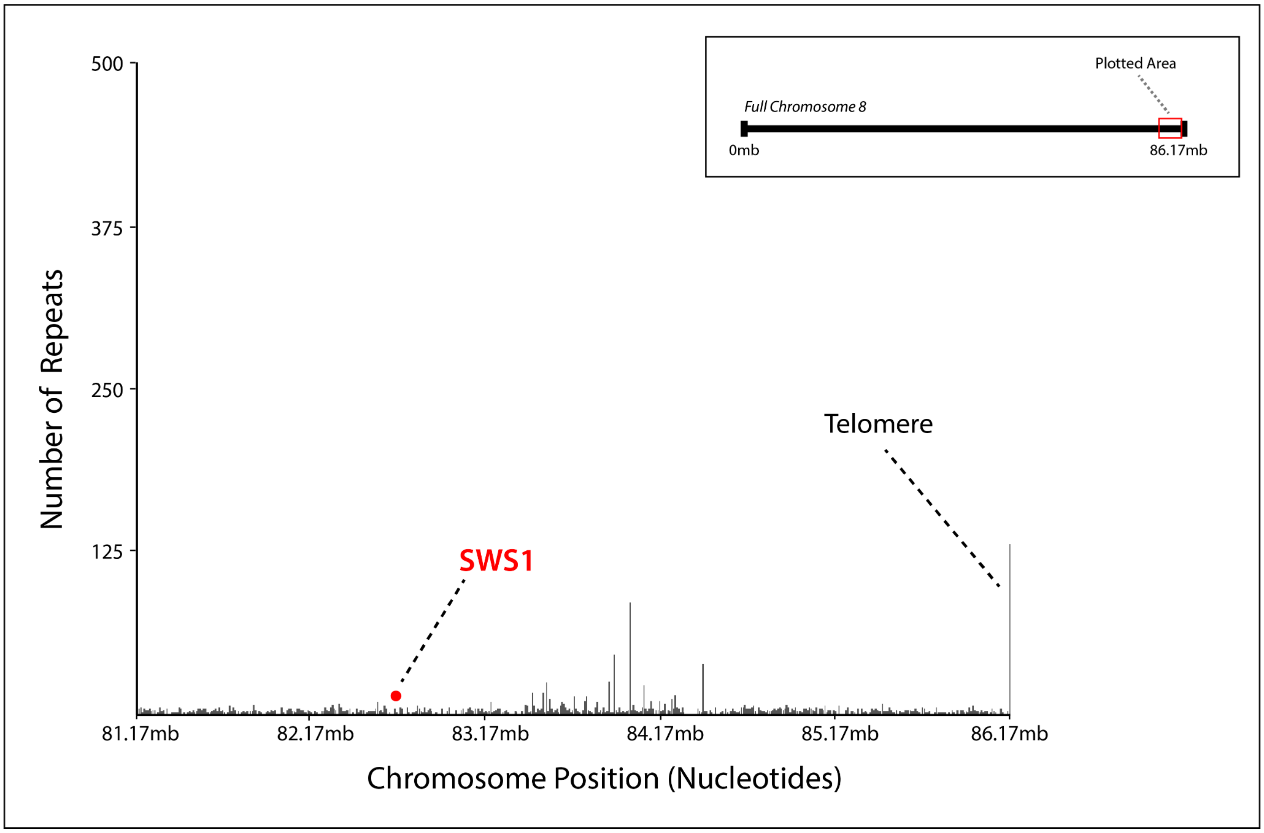


***Figure.S10: Telomere proximity to the SWS1 gene on chromosome 8 of the B. multicinctus genome.*** *The y-axis displays the number of TTAGGG telomere repeat sequences present at a particular contig position on the x-axis. Sequence obtained from Xu et al. 2023 (Genbank = CM042848.1).*

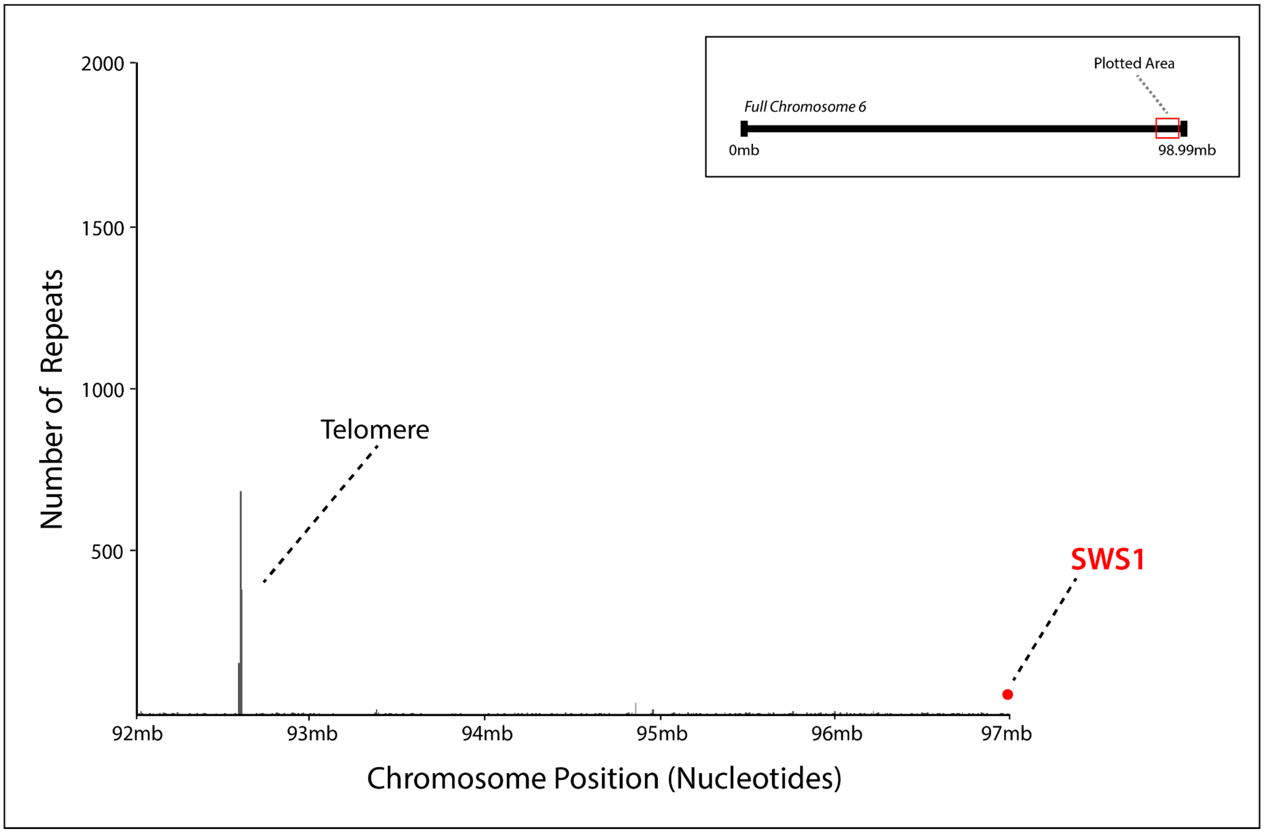


***Figure.S11: Telomere proximity to the SWS1 gene on chromosome 6 of the E. reginae genome.*** *The y-axis displays the number of TTAGGG telomere repeat sequences present at a particular contig position on the x-axis. Sequence obtained from Genbank = CM062046.1.*

### References


Li A, Wang J, Sun K, Wang S, Zhao X, Wang T, Xiong L, Xu W, Qiu L, Shang Y, et al. 2021. Two Reference-Quality Sea Snake Genomes Reveal Their Divergent Evolution of Adaptive Traits and Venom Systems. Molecular Biology and Evolution 38:4867-4883.

Ludington AJ, Hammond JM, Breen J, Deveson IW, Sanders KL. In review. New chromosome-scale genomes provide insights into marine adaptations of sea snakes (Hydrophis: Elapidae). BMC Biol.

Stamatakis A. 2014. RAxML version 8: a tool for phylogenetic analysis and post-analysis of large phylogenies. Bioinformatics 30:1312-1313.

Xu J, Guo S, Yin X, Li M, Su H, Liao X, Li Q, Le L, Chen S, Liao B, et al. 2023. Genomic, transcriptomic, and epigenomic analysis of a medicinal snake, *Bungarus multicinctus*, provides insights into the origin of Elapidae neurotoxins. Acta Pharmaceutica Sinica B 13:2234-2249.
