## Supplementary Tables for "Dynamic Expansions and Retinal Expression of Spectrally Distinct Short-Wavelength Opsin Genes in Sea Snakes"

**Table.S1: Samples used in study.** BioProject = PRJNA1130540.

| Sample ID | Species | Location | Sample Source | Accession # | SWS1 Spectral Tuning Site 86 Genotype |
|---|---|---|---|---|---|
| **Sanger Sequencing of SWS1 Exon 1** | | | | | |
| A00998 | *Hydrophis cyanocinctus* | Gulf of Carpentaria, Australia | Data from Simões et al. 2020, DOI: https://doi.org/10.1016/j.cub.2020.04.061 | MT337758 | Y |
| R37028 | *Hydrophis cyanocinctus* | Gulf of Carpentaria, Australia | Provided by by MAGNT | PP486423 | FY |
| FMNH234200 | *Hydrophis cyanocinctus* | Sabah | Data from Simões et al. 2020, DOI: https://doi.org/10.1016/j.cub.2020.04.061 | MT337911 | Y |
| FMNH234206 | *Hydrophis cyanocinctus* | Sabah | Data from Simões et al. 2020, DOI: https://doi.org/10.1016/j.cub.2020.04.062 | MT337912 | Y |
| FMNH234213 | *Hydrophis cyanocinctus* | Sabah | Data from Simões et al. 2020, DOI: https://doi.org/10.1016/j.cub.2020.04.063 | MT337913 | Y |
| FMNH234219 | *Hydrophis cyanocinctus* | Sabah | Data from Simões et al. 2020, DOI: https://doi.org/10.1016/j.cub.2020.04.064 | MT337914 | Y |
| FMNH249606 | *Hydrophis cyanocinctus* | Phuket | Data from Simões et al. 2020, DOI: https://doi.org/10.1016/j.cub.2020.04.065 | MT337915 | Y |
| FMNH249604 | *Hydrophis cyanocinctus* | Phuket | Data from Simões et al. 2020, DOI: https://doi.org/10.1016/j.cub.2020.04.066 | MT337917 | FY |
| GS140 | *Hydrophis cyanocinctus* | East Java | Data from Simões et al. 2020, DOI: https://doi.org/10.1016/j.cub.2020.04.067 | MT337916 | FY |
| MW04633 | *Hydrophis cyanocinctus* | West Java | Data from Simões et al. 2020, DOI: https://doi.org/10.1016/j.cub.2020.04.068 | MT337918 | F |
| MW04635 | *Hydrophis cyanocinctus* | West Java | Data from Simões et al. 2020, DOI: https://doi.org/10.1016/j.cub.2020.04.069 | MT337919 | F |
| KLS0036 | *Hydrophis cyanocinctus* | Sri Lanka | Data from Simões et al. 2020, DOI: https://doi.org/10.1016/j.cub.2020.04.070 | MT337920 | Y |
| KLS0037 | *Hydrophis cyanocinctus* | Sri Lanka | Data from Simões et al. 2020, DOI: https://doi.org/10.1016/j.cub.2020.04.071 | MT337921 | Y |
| KLS0061 | *Hydrophis cyanocinctus* | Sri Lanka | Data from Simões et al. 2020, DOI: https://doi.org/10.1016/j.cub.2020.04.072 | MT337922 | FY |
| J81639 | *Hydrophis cyanocinctus* | Gulf of Carpentaria, Australia | Provided by by MAGNT | PP486424 | FY |
| J81213 | *Hydrophis cyanocinctus* | Gulf of Carpentaria, Australia | Provided by by MAGNT | PP486425 | FY |
| R36501 | *Hydrophis cyanocinctus* | Gulf of Carpentaria, Australia | Provided by by MAGNT | PP486426 | FY |
| ARR-004 | *Hydrophis cyanocinctus* | Gulf of Thailand | Purchased from Vietnam 2001 | PP486427 | FY |
| ss_UAE598 | *Hydrophis cyanocinctus* | Gulf of Oman | Ludington et al. 2023, sample provided by Belazs Buzas | PP486428 | FY |
| ARR-005 | *Hydrophis cyanocinctus* | Gulf of Thailand | Purchased from Vietnam 2001 | PP486429 | FY |
| ARR-003 | *Hydrophis cyanocinctus* | Gulf of Thailand | Purchased from Vietnam 2001 | PP486430 | F |
| A022 | *Hydrophis cyanocinctus* | Gulf of Thailand | Purchased from Vietnam 2001 | PP486431 | F |
| 260496 | *Hydrophis cyanocinctus* | Thailand | Provided by FMNH | PP486432 | F |
| 249657 | *Hydrophis cyanocinctus* | Thailand | Provided by FMNH | PP486433 | FY |
| 249612 | *Hydrophis cyanocinctus* | Phuket, Thailand | Provided by FMNH | PP486434 | FY |
| 249609 | *Hydrophis cyanocinctus* | Phuket, Thailand | Provided by FMNH | PP486435 | F |
| cyn4 | *Hydrophis cyanocinctus* | Gulf of Thailand | Purchased from Vietnam 2019 | PP486436 | FY |
| MW04636 | *Hydrophis cyanocinctus* | West Java, Indonesia | Provided by Bogor Museum, Indonesia | PP486437 | FY |
| MW04634 | *Hydrophis cyanocinctus* | West Java, Indonesia | Provided by Bogor Museum, Indonesia | PP486438 | F |
| ARR-006 | *Hydrophis cyanocinctus* | Gulf of Thailand | Purchased from Vietnam 2001 | PP486439 | FY |
| ARR-002 | *Hydrophis cyanocinctus* | Gulf of Thailand | Purchased from Vietnam 2001 | PP486440 | FY |
| R37129 | *Hydrophis cyanocinctus* | Gulf of Carpentaria, Australia | Provided by by MAGNT | PP486441 | FY |
| R37126 | *Hydrophis cyanocinctus* | Gulf of Carpentaria, Australia | Provided by by MAGNT | PP486442 | FY |
| J81022 | *Hydrophis cyanocinctus* | Gulf of Carpentaria, Australia | Provided by by MAGNT | PP486443 | FY |
| R37125 | *Hydrophis cyanocinctus* | Gulf of Carpentaria, Australia | Provided by by MAGNT | PP486444 | FY |
| cyano1998/AK39 | *Hydrophis cyanocinctus* | Phuket, Thailand | Provided by FMNH | PP486445 | FY |
| 260497 | *Hydrophis cyanocinctus* | Thailand | Provided by FMNH | PP486446 | Y |
| 249610 | *Hydrophis cyanocinctus* | Phuket, Thailand | Provided by FMNH | PP486447 | FY |
| 249562 | *Hydrophis cyanocinctus* | Phuket, Thailand | Provided by FMNH | PP486448 | Y |
| 249525 | *Hydrophis cyanocinctus* | Phuket, Thailand | Provided by FMNH | PP486449 | Y |
| 249519 | *Hydrophis cyanocinctus* | Phuket, Thailand | Provided by FMNH | PP486450 | Y |
| 249517 | *Hydrophis cyanocinctus* | Phuket, Thailand | Provided by FMNH | PP486451 | FY |
| 249490 | *Hydrophis cyanocinctus* | Phuket, Thailand | Provided by FMNH | PP486452 | FY |
| 234208 | *Hydrophis cyanocinctus* | East Malaysia | Provided by FMNH | PP486453 | Y |
| 234202 | *Hydrophis cyanocinctus* | East Malaysia | Provided by FMNH | PP486454 | Y |
| 234201 | *Hydrophis cyanocinctus* | East Malaysia | Provided by FMNH | PP486455 | Y |
| cyn5 | *Hydrophis cyanocinctus* | Gulf of Thailand | Purchased from Vietnam 2019 | PP486456 | FY |
| cyn3 | *Hydrophis cyanocinctus* | Gulf of Thailand | Purchased from Vietnam 2019 | PP486457 | Y |
| cyn2 | *Hydrophis cyanocinctus* | Gulf of Thailand | Purchased from Vietnam 2019 | PP486458 | Y |
| cyn1 | *Hydrophis cyanocinctus* | Gulf of Thailand | Purchased from Vietnam 2019 | PP486459 | FY |
| 249460 | *Hydrophis cyanocinctus* | Phuket, Thailand | Provided by FMNH | PP486460 | FY |
| 234182 | *Hydrophis cyanocinctus* | East Malaysia | Provided by FMNH | PP486461 | Y |
| ss_UAE742 | *Hydrophis cyanocinctus* | Gulf of Oman | Ludington et al. 2023, sample provided by Belazs Buzas | PP486462 | FY |
| KLS1070 | *Hydrophis fasciatus* | Penang, Malaysia | Provided by Luke Allen | PP624366 | FY |
| KLS1071 | *Hydrophis fasciatus* | Penang, Malaysia | Provided by Luke Allen | PP624393 | FY |
| KLS1074 | *Hydrophis fasciatus* | Penang, Malaysia | Provided by Luke Allen | PP624365 | FY |
| KLS1076 | *Hydrophis fasciatus* | Penang, Malaysia | Provided by Luke Allen | PP624364 | FY |
| KLS1077 | *Hydrophis fasciatus* | Penang, Malaysia | Provided by Luke Allen | PP624371 | FY |
| KLS1078 | *Hydrophis fasciatus* | Penang, Malaysia | Provided by Luke Allen | PP624376 | FY |
| KLS1080 | *Hydrophis fasciatus* | Penang, Malaysia | Provided by Luke Allen | PP624381 | FY |
| KLS1081 | *Hydrophis fasciatus* | Penang, Malaysia | Provided by Luke Allen | PP624394 | FY |
| X01002 | *Hydrophis atriceps* | Gulf of Thailand | Data from Simões et al. 2020, DOI: https://doi.org/10.1016/j.cub.2020.04.061 | MT337886 | FY |
| X01003 | *Hydrophis atriceps* | Gulf of Thailand | Data from Simões et al. 2020, DOI: https://doi.org/10.1016/j.cub.2020.04.061 | MT337887 | Y |
| X01022 | *Hydrophis atriceps* | Gulf of Thailand | Data from Simões et al. 2020, DOI: https://doi.org/10.1016/j.cub.2020.04.061 | MT337888 | FY |
| atr1_VT | *Hydrophis atriceps* | Gulf of Thailand | Purchased from Vietnam 2019 | PP624372 | FY |
| atr2_VT/VT2 | *Hydrophis atriceps* | Gulf of Thailand | Purchased from Vietnam 2019 | PP624391 | FY |
| atr3_VT | *Hydrophis atriceps* | Gulf of Thailand | Purchased from Vietnam 2019 | PP624374 | FY |
| atr4_VT | *Hydrophis atriceps* | Gulf of Thailand | Purchased from Vietnam 2019 | PP624389 | FY |
| atr5_VT | *Hydrophis atriceps* | Gulf of Thailand | Purchased from Vietnam 2019 | PP624375 | FY |
| atr7_VT | *Hydrophis atriceps* | Gulf of Thailand | Purchased from Vietnam 2019 | PP624377 | FY |
| atr8_VT | *Hydrophis atriceps* | Gulf of Thailand | Purchased from Vietnam 2019 | PP624387 | FY |
| fas1_VT | *Hydrophis atriceps* | Gulf of Thailand | Purchased from Vietnam 2019 | PP624369 | FY |
| fas2_VT | *Hydrophis atriceps* | Gulf of Thailand | Purchased from Vietnam 2019 | PP624368 | FY |
| par1_VT | *Hydrophis atriceps* | Gulf of Thailand | Purchased from Vietnam 2019 | PP624386 | FY |
| MW04643 | *Hydrophis spiralis* | West Java, Indonesia | Data from Simões et al. 2020, DOI: https://doi.org/10.1016/j.cub.2020.04.061 | MT337977 | F |
| MW04646 | *Hydrophis spiralis* | West Java, Indonesia | Provided by Bogor Museum, Indonesia | PP624398 | FY |
| MW04650 | *Hydrophis spiralis* | West Java, Indonesia | Provided by Bogor Museum, Indonesia | PP624383 | FY |
| MW04660 | *Hydrophis spiralis* | West Java, Indonesia | Provided by Bogor Museum, Indonesia | PP624382 | FY |
| MW04662 | *Hydrophis spiralis* | West Java, Indonesia | Provided by Bogor Museum, Indonesia | PP624403 | FY |
| MW04663 | *Hydrophis spiralis* | West Java, Indonesia | Provided by Bogor Museum, Indonesia | PP624396 | FY |
| MW04651 | *Hydrophis spiralis* | West Java, Indonesia | Provided by Bogor Museum, Indonesia | PP624404 | FY |
| MW04652 | *Hydrophis spiralis* | West Java, Indonesia | Provided by Bogor Museum, Indonesia | PP624402 | FY |
| MW04653 | *Hydrophis spiralis* | West Java, Indonesia | Provided by Bogor Museum, Indonesia | PP624379 | FY |
| MW04654 | *Hydrophis spiralis* | West Java, Indonesia | Provided by Bogor Museum, Indonesia | PP624400 | F |
| MW04655 | *Hydrophis spiralis* | West Java, Indonesia | Provided by Bogor Museum, Indonesia | PP624380 | F |
| MW04657 | *Hydrophis spiralis* | West Java, Indonesia | Provided by Bogor Museum, Indonesia | PP624384 | FY |
| MW04658 | *Hydrophis spiralis* | West Java, Indonesia | Provided by Bogor Museum, Indonesia | PP624397 | F |
| MW04664 | *Hydrophis spiralis* | West Java, Indonesia | Provided by Bogor Museum, Indonesia | PP624401 | FY |
| X01010 | *Hydrophis parviceps* | Vietnam | Data from Simões et al. 2020, DOI: https://doi.org/10.1016/j.cub.2020.04.061 | MT337766 | F |
| X01011 | *Hydrophis parviceps* | Vietnam | Data from Simões et al. 2020, DOI: https://doi.org/10.1016/j.cub.2020.04.061 | MT337967 | F |
| X01012 | *Hydrophis parviceps* | Vietnam | Data from Simões et al. 2020, DOI: https://doi.org/10.1016/j.cub.2020.04.061 | MT337968 | Y |
| MT189 | *Hydrophis coggeri* | South Sulawesi Selatan | Data from Simões et al. 2020, DOI: https://doi.org/10.1016/j.cub.2020.04.061 | MT337900 | F |
| MT190 | *Hydrophis coggeri* | South Sulawesi Selatan | Data from Simões et al. 2020, DOI: https://doi.org/10.1016/j.cub.2020.04.061 | MT337901 | F |
| MT162 | *Hydrophis coggeri* | South Sulawesi Selatan | Provided by Bogor Museum, Indonesia | PP624373 | F |
| MT171 | *Hydrophis coggeri* | South Sulawesi Selatan | Provided by Bogor Museum, Indonesia | PP624388 | F |
| MT172 | *Hydrophis coggeri* | South Sulawesi Selatan | Provided by Bogor Museum, Indonesia | PP624385 | F |
| MT174 | *Hydrophis coggeri* | South Sulawesi Selatan | Provided by Bogor Museum, Indonesia | PP624390 | F |
| MT179 | *Hydrophis coggeri* | South Sulawesi Selatan | Provided by Bogor Museum, Indonesia | PP624392 | F |
| MW04740 | *Hydrophis coggeri* | South Sulawesi Selatan | Provided by Bogor Museum, Indonesia | PP624399 | FY |
| MW04741 | *Hydrophis coggeri* | South Sulawesi Selatan | Provided by Bogor Museum, Indonesia | PP624378 | FY |
| MW04743 | *Hydrophis coggeri* | South Sulawesi Selatan | Provided by Bogor Museum, Indonesia | PP624395 | FY |
| mel1_VT | *Hydrophis belcheri* | Gulf of Thailand | Purchased from Vietnam 2019 | PP624370 | Y |
| mel2_VT | *Hydrophis belcheri* | Gulf of Thailand | Purchased from Vietnam 2019 | PP624367 | FY |
| X01008 | *Hydrophis belcheri* | Gulf of Thailand | Data from Simões et al. 2020, DOI: https://doi.org/10.1016/j.cub.2020.04.061 | MT337889 | Y |
| **Transcriptome Analysis** | | | | | |
| atr3_VT | *Hydrophis atriceps* | Gulf of Thailand | Purchased from Vietnam 2019 | SAMN42208552 | |
| atr4_VT | *Hydrophis atriceps* | Gulf of Thailand | Purchased from Vietnam 2019 | SAMN42208553 | |
| cyn4_VT | *Hydrophis cyanocinctus* | Gulf of Thailand | Purchased from Vietnam 2019 | SAMN42208554 | |

| | | | | |
|---|---|---|---|---|
| cyn5_VT | *Hydrophis cyanocinctus* | Gulf of Thailand | Purchased from Vietnam 2019 | SAMN42208555 |
| KLS0587 | *Aipysurus mosaicus* | Broome, Australia | Field expedition 2014 | SAMN42208561 |
| KLS0459 | *Aipysurus laevis* | Broome, Australia | Field expedition 2014 | SAMN42208556 |
| KLS0517 | *Hydrophis peronii* | Broome, Australia | Field expedition 2014 | SAMN42208558 |
| KLS0515 | *Hydrophis kingii* | Broome, Australia | Field expedition 2014 | SAMN42208557 |
| KLS0534 | *Hydrelaps darwiniensis* | Broome, Australia | Field expedition 2014 | SAMN42208559 |
| KLS0560 | *Hydrophis major* | Broome, Australia | Field expedition 2014 | SAMN42208560 |
| **Long-Read Whole Genomes** | | | | |
| QH_4_m | *Hydrophis cyanocinctus* | South China Sea | Data from Li et al. 2021, DOI: msab212 | JAAZTL000000000.1 |
| PK_4_m | *Hydophis curtus* | South China Sea | Data from Li et al. 2021, DOI: msab212 | JABAHG000000000.1 |
| ss_UAE728 | *Hydrophis curtus* | Gulf of Oman | Ludington et al. 2023, sample provided by Belazs Buzas | SAMN23222007 |
| KLS1140 | *Aipysurus laevis* | Broome, Australia | Field expedition 2021 | PRJNA1115802 |
| KLS1130 | *Hydrophis major* | Broome, Australia | Ludington et al. 2023, field expedition 2021 | SAMN35768000 |
| KLS1121 | *Hydrophis elegans* | Broome, Australia | Ludington et al. 2023, field expedition 2021 | SAMN35787919 |
| nn02 | *Naja naja* | Kerala, India | Suryamohan et al. 2020 | GCA_009733165 |
| TS10Xv2-PRI | *Notechis scutatus* | Australia | Edwards et al. 2018, DOI: f1000research.1115550.1 | GCA_900518725.1 |
| SS_UAE730 | *Hydrophis ornatus* | Gulf of Oman | Ludington et al. 2023, sample provided by Belazs Buzas | SAMN23222008 |
| **Short-Read Genome Coverage Analysis** | | | | |
| KLS1272 | *Hydrelaps darwiniensis* | Broome, Australia | Field expedition 2022 | SAMN42208587 |
| atr2_VT/VTA2 | *Hydrophis atriceps* | Gulf of Thailand | Purchased from Vietnam 2019 | SAMN42208566 |
| cur1VT | *Hydrophis curtus* | Gulf of Thailand | Purchased from Vietnam 2019 | SAMN42208567 |
| KLS1024 | *Hydrophis elegans* | Broome, Australia | Field expedition 2018 | SAMN42208582 |
| KLS1140 | *Aipysurus laevis* | Broome, Australia | Field expedition 2021 | PRJNA1115802 |
| KLS1167 | *Aipysurus laevis* | Broome, Australia | Field expedition 2021 | SAMN42208586 |
| KLS0691 | *Aipysurus laevis* | Broome, Australia | Field expedition 2016 | PRJNA669384 |
| cur2VT | *Hydrophis curtus* | Gulf of Thailand | Purchased from Vietnam 2019 | SAMN42208568 |
| cur3VT | *Hydrophis curtus* | Gulf of Thailand | Purchased from Vietnam 2019 | SAMN42208569 |
| cur4VT | *Hydrophis curtus* | Gulf of Thailand | Purchased from Vietnam 2019 | SAMN42208570 |
| cur7VT | *Hydrophis curtus* | Gulf of Thailand | Purchased from Vietnam 2019 | SAMN42208571 |
| R36639 | *Hydrophis curtus* | Gulf of Thailand | Purchased from Vietnam 2019 | SAMN42208600 |
| KLS1025 | *Hydrophis elegans* | Broome, Australia | Field expedition 2018 | SAMN42208583 |
| KLS1027 | *Hydrophis elegans* | Broome, Australia | Field expedition 2018 | SAMN42208584 |
| KLS1034 | *Hydrophis elegans* | Broome, Australia | Field expedition 2018 | SAMN42208585 |
| KLS1121 | *Hydrophis elegans* | Broome, Australia | Field expedition 2021 | SAMN35787919 |
| KLS0629 | *Hydrophis major* | Broome, Australia | Field expedition 2015 | SAMN42208575 |
| KLS0630 | *Hydrophis major* | Broome, Australia | Field expedition 2015 | SAMN42208576 |
| KLS0631 | *Hydrophis major* | Broome, Australia | Field expedition 2015 | SAMN42208577 |
| KLS0647 | *Hydrophis major* | Broome, Australia | Field expedition 2015 | SAMN42208578 |
| KLS1020 | *Hydrophis major* | Broome, Australia | Field expedition 2018 | SAMN42208581 |
| KLS1018 | *Hydrophis ocellatus* | Broome, Australia | Field expedition 2018 | SAMN42208580 |
| FMNH249563 | *Hydrophis ornatus* | Phuket Province, Thailand | Provided by FMNH | SAMN42208572 |
| KLS1292 | *Hydrophis peronii* | Pilbara, Australia | Field expedition 2022 | SAMN42208589 |
| KLS0095 | *Hydrophis platurus* | Mannar, Sri Lanka | Provided by Kanishka Ukuwela | SAMN42208574 |
| KLS0026 | *Hydrophis schistosus* | Puttlam, Sri Lanka | Provided by Kanishka Ukuwela | SAMN42208573 |
| LAHSC07 | *Hydrophis schistosus* | Penang, Malaysia | Provided by Luke Allen | SAMN42208591 |
| KLS0938 | *Hydrophis stokesii* | Exmouth, Australia | Field expedition 2017 | SAMN42208579 |
| nn01 | *Naja naja* | Kerala, India | Suryamohan et al. 2020 | GCA_009733165 |
| nn02 | *Naja naja* | Kerala, India | Suryamohan et al. 2020 | GCA_009733165 |
| TS10Xv2-PRI | *Notechis scutatus* | Australia | Edwards et al. 2018, DOI: f1000research.1115550.1 | GCA_900518725.1 |
| ARR-002 | *Hydrophis cyanocinctus* | Gulf of Thailand | Purchased from Vietnam 2001, sequenced by Vhon Garcia | SAMN42208562 |
| ARR-003 | *Hydrophis cyanocinctus* | Gulf of Thailand | Purchased from Vietnam 2001, sequenced by Vhon Garcia | SAMN42208563 |
| ARR-004 | *Hydrophis cyanocinctus* | Gulf of Thailand | Purchased from Vietnam 2001, sequenced by Vhon Garcia | SAMN42208564 |
| ARR-005 | *Hydrophis cyanocinctus* | Gulf of Thailand | Purchased from Vietnam 2001, sequenced by Vhon Garcia | SAMN42208565 |
| SD_cyn2 | *Hydrophis cyanocinctus* | Gulf of Thailand | Purchased from Vietnam 2019, sequenced by Vhon Garcia | SAMN42208601 |
| SD_cyn3 | *Hydrophis cyanocinctus* | Gulf of Thailand | Purchased from Vietnam 2019, sequenced by Vhon Garcia | SAMN42208602 |
| VT_cyn1 | *Hydrophis cyanocinctus* | Gulf of Thailand | Purchased from Vietnam 2019, sequenced by Vhon Garcia | SAMN42208604 |
| VT_cyn2 | *Hydrophis cyanocinctus* | Gulf of Thailand | Purchased from Vietnam 2019, sequenced by Vhon Garcia | SAMN42208605 |
| SD_mel1 | *Hydrophis melanocephalus* | Gulf of Thailand | Purchased from Vietnam 2019, sequenced by Vhon Garcia | SAMN42208603 |
| MT164 | *Hydrophis melanocephalus* | Makassar, Indonesia | Provided by Bogor Museum, Indonesia; sequenced by Vhon Garcia | SAMN42208593 |
| MT167 | *Hydrophis melanocephalus* | Makassar, Indonesia | Provided by Bogor Museum, Indonesia; sequenced by Vhon Garcia | SAMN42208594 |
| MT169 | *Hydrophis melanocephalus* | Makassar, Indonesia | Provided by Bogor Museum, Indonesia; sequenced by Vhon Garcia | SAMN42208595 |
| MT185 | *Hydrophis melanocephalus* | Makassar, Indonesia | Provided by Bogor Museum, Indonesia; sequenced by Vhon Garcia | SAMN42208596 |
| MT186 | *Hydrophis melanocephalus* | Makassar, Indonesia | Provided by Bogor Museum, Indonesia; sequenced by Vhon Garcia | SAMN42208597 |
| MT187 | *Hydrophis melanocephalus* | Makassar, Indonesia | Provided by Bogor Museum, Indonesia; sequenced by Vhon Garcia | SAMN42208598 |
| MW04737 | *Hydrophis melanocephalus* | Makassar, Indonesia | Provided by Bogor Museum, Indonesia; sequenced by Vhon Garcia | SAMN42208599 |
| MT04738 | *Hydrophis melanocephalus* | Makassar, Indonesia | Provided by Bogor Museum, Indonesia; sequenced by Vhon Garcia | SAMN42208592 |
| KLS1291 | *Hydrophis kingii* | Broome, Australia | Field expedition 2022 | SAMN42208588 |
| KLS1318 | *Hydrophis spiralis* | Gulf of Thailand | Purchased from Vietnam 2022 | SAMN42208590 |
| **Chromosome Annotation of Terrestrial Snake Outgroups** | | | | |
| - | *Ahaetulla prasina* | Xishuangbanna, China | Tang et al. 2023 | NC_080545.1 |
| - | *Bungarus multicinctus* | China | Xu et al. 2023 | CM042848.1 |
| - | *Erythrolamprus reginae* | Reserva Natural Tanimboca, Colombia | Tarvin et al. (unpublished) | CM062046.1 |
| - | *Liasis olivaceus* | Perth, Australia | Allentoft et al. (unpublished) | CM061355.1 |
| - | *Vipera latastei* | Beceite, Spain | Marques-Bonet et al. (unpublished) | CM044359.1 |

**References**


Edwards R, Amos T, Tang J. 2018. Pseudodiploid pseudo-long-read whole genome sequencing and assembly of Pseudonaja textilis (eastern brown snake) and Notechis scutatus (mainland tiger snake). In: F1000Research. p. 753.
Li A, Wang J, Sun K, Wang S, Zhao X, Wang T, Xiong L, Xu W, Qiu L, Shang Y, et al. 2021. Two Reference-Quality Sea Snake Genomes Reveal Their Divergent Evolution of Adaptive Traits and Venom Systems. Molecular Biology and Evolution 38:4867-4883.
Ludington AJ, Hammond JM, Breen J, Deveson IW, Sanders KL. In review. New chromosome-scale genomes provide insights into marine adaptations of sea snakes (Hydrophis: Elapidae). BMC Biol.
Simões BF, Gower DJ, Rasmussen AR, Sarker MAR, Fry GC, Casewell NR, Harrison RA, Hart NS, Partridge JC, Hunt DM, et al. 2020. Spectral Diversification and Trans-Species Allelic Polymorphism during the Land-to-Sea Transition in Snakes. Current Biology.
Suryamohan K, Krishnankutty SP, Guillory J, Jevit M, Schröder MS, Wu M, Kuriakose B, Mathew OK, Perumal RC, Koludarov I, et al. 2020. The Indian cobra reference genome and transcriptome enables comprehensive identification of venom toxins. Nature Genetics 52: 106-117.
Tang C-Y, Zhang X, Xu X, Sun S, Peng C, Song M-H, Yan C, Sun H, Liu M, Xie L, et al. 2023. Genetic mapping and molecular mechanism behind color variation in the Asian vine snake. Genome Biology 24:46.
Xu J, Guo S, Yin X, Li M, Su H, Liao X, Li Q, Le L, Chen S, Liao B, et al. 2023. Genomic, transcriptomic, and epigenomic analysis of a medicinal snake, Bungarus multicinctus, to provides insights into the origin of Elapidae neurotoxins. Acta Pharmaceutica Sinica B 13:2234-2249.

**Table.S2:** ***Aipysurus laevis*** **genome assembly quality information.**

| K-mer completeness | | | | |
|---|---|---|---|---|
| **Genome** | **K-mer set** | **Solid k-mers (asm)** | **Solid k-mers (reads)** | **Completeness (%)** |
| *Aipysurus laevis* | all | 1,121,717,406 | 1,196,647,986 | 93.7 |

| Consensus Quality (QV) | | | | |
|---|---|---|---|---|
| **Genome** | **Unique k-mers (asm)** | **Unique k-mers (reads)** | **QV** | **Error rate** |
| *Aipysurus laevis* | 23,645,419 | 2,076,720,583 | 32.6 | 0.000545149 |

| **Assembly** | |
|---|---|
| # contigs (>= 0 bp) | 4371 |
| # contigs (>= 1000 bp) | 4297 |
| # contigs (>= 5000 bp) | 3690 |
| # contigs (>= 10000 bp) | 2712 |
| # contigs (>= 25000 bp) | 1435 |
| # contigs (>= 50000 bp) | 1017 |
| Total length (>= 0 bp) | 2076908980 |
| Total length (>= 1000 bp) | 2076857158 |
| Total length (>= 5000 bp) | 2074803903 |
| Total length (>= 10000 bp) | 2067599300 |
| Total length (>= 25000 bp) | 2047919121 |
| Total length (>= 50000 bp) | 2033393357 |
| # contigs | 4370 |
| Largest contig | 99471500 |
| Total length | 2076908564 |
| GC (%) | 40.92 |
| N50 | 27079146 |
| N75 | 9157208 |
| L50 | 21 |
| L75 | 52 |
| # N's per 100 kbp | 0.27 |

| **Flow Cell Statistics** | **PBXP22284 (flow cell 1)** | **PUXP22260 (flow cell 2)** | **Total** |
|---|---|---|---|
| Number of reads | 2,846,372 | 1,532,417 | 4,378,789 |
| Number of bases | 49,580,515,741 | 37,578,309,092 | 87,158,824,833 |
| N50 read length | 34,805 | 52,591 | 40,299 |
| Longest read | 401,509 | 524,290 | 524,290 |
| Shortest read | 12 | 16 | 12 |
| Mean read length | 17,418 | 24,522 | 19,904 |
| Median read length | 9,820 | 12,800 | 10,813 |
| Mean read quality | 14.14 | 14.53 | 14.35 |
| Median read quality | 14.6 | 14.81 | 14.68 |

**Table.S3: SWS1 information for snake genomes assembled from long-read data.**

Note that one SWS1 copy in *H. cyanocinctus* is suspected to be a product of assembly error, as detailed in the main text. However, the false copy cannot be determined at this time, so all are included in the below table.

| Species | Chromosome/ Scaffold | Notes | Start Position (nt) | End Position (nt) | Targeted Coverage |
|---|---|---|---|---|---|
| *Hydrophis cyanocinctus* | Chromosome 4 | Copy A | 18,517 | 30,599 | 70x PacBio |
| *Hydrophis cyanocinctus* | Chromosome 4 | Copy B | 62,265 | 74,391 | 70x PacBio |
| *Hydrophis cyanocinctus* | Chromosome 4 | Copy C (inverted) | 109,145 | 121,468 | 70x PacBio |
| *Hydrophis cyanocinctus* | Chromosome 4 | Copy D (inverted) | 3,832,469 | 3,844,440 | 70x PacBio |
| *Hydrophis curtus* (Li et al. 2021) | Chromosome 4 | (inverted) | 216,835,270 | 216,847,357 | 70x PacBio |
| *Hydrophis curtus* (Ludington et al. 2023) | Chromosome 4 | - | 2,478,869 | 2,490,329 | 60x Nanopore |
| *Aipysurus laevis* | Contig_95 | - | 2,698,255 | 2,711,527 | 40x Nanopore |
| *Hydrophis major* | Chromosome 4 | (inverted) | 213,702,160 | 213,714,105 | 30x HiFi |
| *Hydrophis elegans* | Contig_3820 | - | 243,733 | 255,644 | 40x Nanopore |
| *Hydrophis ornatus* | Chromosome 4 | (inverted) | 212,420,261 | 212,421,219 | 60x Nanopore |
| *Naja naja* | Chromosome 2 (exon 1) | - | 456 | 987 | 60x Nanopore |
| *Notechis scutatus* | ULFQ01013886 (exon 1) | - | 2,701 | 3,066 | 71x PacBio |

| Table.S4: Variant frequency at spectral tuning site 86. | | | | |
|---|---|---|---|---|
| | | Number of Reads at Site 86 | | |
| **Species** | **Sample ID** | **F86** | **Y86** | **S86** |
| *Aipysurus laevis* | KLS0691 | 0 | 0 | 34 |
| | KLS1140 | 1 | 0 | 53 |
| | KLS1167 | 1 | 0 | 21 |
| *Hydrelaps darwiniensis* | KLS1272 | 24 | 0 | 0 |
| *Hydrophis atriceps* | VTA_2 | 24 | 24 | 0 |
| *Hydrophis curtus* | cur1VT | 2 | 22 | 0 |
| | cur2VT | 1 | 20 | 0 |
| | cur3VT | 13 | 16 | 0 |
| | cur4VT | 0 | 30 | 0 |
| | cur7VT | 0 | 19 | 0 |
| | R36639 | 0 | 28 | 1 |
| *Hydrophis cyanocinctus* | ARR_002 | 5 | 2 | 0 |
| | ARR_003 | 9 | 0 | 0 |
| | ARR_004 | 2 | 1 | 0 |
| | ARR_005 | 4 | 0 | 0 |
| | SD_cyn2 | 0 | 9 | 0 |
| | SD_cyn3 | 0 | 12 | 0 |
| | VT_cyn1 | 4 | 1 | 0 |
| | VT_cyn2 | 2 | 6 | 0 |
| *Hydrophis elegans* | KLS1024 | 20 | 1 | 0 |
| | KLS1025 | 20 | 0 | 0 |
| | KLS1027 | 23 | 0 | 0 |
| | KLS1034 | 27 | 0 | 0 |
| | KLS1121 | 68 | 1 | 0 |
| *Hydrophis kingii* | KLS1291 | 0 | 101 | 0 |
| *Hydrophis major* | KLS0629 | 15 | 0 | 0 |
| | KLS9630 | 24 | 0 | 0 |
| | KLS0631 | 21 | 0 | 0 |
| | KLS0647 | 14 | 0 | 0 |
| | KLS1020 | 18 | 0 | 0 |
| *Hydrophis melanocephalus* | SD_mel1 | 2 | 2 | 0 |
| | MT164 | 2 | 0 | 0 |
| | MT167 | 1 | 0 | 0 |
| | MT186 | 4 | 0 | 0 |
| | MW04737 | 4 | 0 | 0 |
| | MT04738 | 3 | 3 | 0 |
| | MT169 | 0 | 1 | 0 |
| | MT185 | 0 | 1 | 0 |
| | MT187 | 1 | 1 | 0 |
| *Hydrophis ocellatus* | KLS1018 | 0 | 27 | 0 |

| | | | | |
|---|---|---|---|---|
| *Hydrophis ornatus* | FMNH249563 | 0 | 9 | 0 |
| *Hydrophis peronii* | KLS1292 | 0 | 24 | 0 |
| *Hydrophis platurus* | KLS0095 | 27 | 0 | 0 |
| *Hydrophis schistosus* | KLS0026 | 101 | 0 | 0 |
| | LAHSc07 | 103 | 0 | 0 |
| *Hydrophis spiralis* | KLS1318 | 40 | 48 | 0 |
| *Hydrophis stokesii* | KLS0938 | 0 | 15 | 0 |

**Table.S5: SWS1 gene position in relation to chromosome sequence end for three terrestrial snakes.**

*Terrestrial snakes with chromosome-level assemblies were selected. Genomes presented here are those which do not have telomeric repeats close to the SWS1 gene (see supplementary figs.S6-11 for those that were in close proximity). The SWS1 genes in A. prasina and L. olivaceous are in close proximity to sequence ends and are therefore likely to be at chromatid ends also. The fact that telomeric repeats do not also appear close in these genomes is likely due to the assembly mis-assigning telomere-containing reads to one end of the sequence only.*

| Species | Chromosome | Chromosome Length | SWS1 Start Position | SWS1 End Position | SWS1 Gene Proximity to Sequence End |
|---|---|---|---|---|---|
| *Ahaetulla prasina* | 7 | 103,430,753 | 100,939,590 | 100,947,758 | 2.4% of full chromosome length |
| *Liasis olivaceous* | 8 | 82,069,438 | 79,554,274 | 79,561,678 | 3.0% of full chromosome length |
| *Vipera latastei* | 6 | 93,683,332 | 48,988,143 | 48,998,265 | 47.69% of full chromosome length |

**Table.S6: Transcriptome results for RNA-sequencing of SWS1 site 86 in 10 elapid snakes.**

*The displayed results only include reads which passed quality filtering.*

| | Number of Reads at Site 86 | | | |
|---|---|---|---|---|
| **Species/Sample** | **A** | **C** | **G** | **T** |
| *Hydrophis atriceps (A3)* | 100 | 0 | 0 | 113 |
| *Hydrophis atriceps (A4)* | 118 | 0 | 0 | 129 |
| *Hydrophis cyanocinctus (C4)* | 489 | 0 | 0 | 250 |
| *Hydrophis cyanocinctus (C5)* | 483 | 0 | 0 | 415 |
| *Aipysurus laevis* | 0 | 1060 | 0 | 0 |
| *Aipysurus mosaicus* | 0 | 553 | 0 | 0 |
| *Hydrophis peronii* | 1615 | 0 | 0 | 0 |
| *Hydrophis kingii* | 1357 | 0 | 0 | 0 |
| *Hydrelaps darwiniensis* | 0 | 0 | 0 | 363 |
| *Hydrophis major* | 0 | 0 | 0 | 790 |

| Table.S7: Depth statistics for short-read data of 50 elapid snake genomes. | | | | | |
|---|---|---|---|---|---|
| **Sample** | **Species** | **Aligned reads (post-filter)** | **Avg. depth** | **Avg. depth (chr 4)** | **Breadth of coverage (chr4)** |
| KLS0691 | *Aipysurus laevis* | 501,676,686 | 28.2 | 32.01 | 79.78 |
| KLS1140 | *Aipysurus laevis* | 505,949,917 | 32.66 | 38.08 | 83.17 |
| KLS1167 | *Aipysurus laevis* | 192,483,259 | 12.59 | 14.51 | 80.66 |
| VT_cyn2 | *Hydrophis cyanocinctus* | 168,361,648 | 7.44 | 8.34 | 86.07 |
| ARR-002 | *Hydrophis cyanocinctus* | 96,422,148 | 4.56 | 5.1 | 82.58 |
| ARR-003 | *Hydrophis cyanocinctus* | 105,515,503 | 5.07 | 5.68 | 83.50 |
| ARR-004 | *Hydrophis cyanocinctus* | 98,054,560 | 4.73 | 5.55 | 83.46 |
| ARR-005 | *Hydrophis cyanocinctus* | 88,903,011 | 4.02 | 4.76 | 80.50 |
| SD_cyn2 | *Hydrophis cyanocinctus* | 190,040,616 | 9.4 | 10.54 | 87.84 |
| SD_cyn3 | *Hydrophis cyanocinctus* | 133,610,696 | 6.59 | 7.7 | 85.86 |
| VT_cyn1 | *Hydrophis cyanocinctus* | 126,788,217 | 5.94 | 6.88 | 84.60 |
| VTA_2 | *Hydrophis atriceps* | 262,217,108 | 17.72 | 19.83 | 91.82 |
| Cur1VT | *Hydrophis curtus* | 204,655,058 | 14.01 | 15.68 | 91.87 |
| Cur2VT | *Hydrophis curtus* | 207,917,136 | 14.17 | 16.41 | 91.73 |
| Cur3VT | *Hydrophis curtus* | 216,398,521 | 14.73 | 17.08 | 91.93 |
| Cur4VT | *Hydrophis curtus* | 276,894,977 | 18.45 | 21.07 | 92.12 |
| Cur7VT | *Hydrophis curtus* | 190,726,398 | 13.01 | 15.13 | 91.73 |
| KLS1024 | *Hydrophis elegans* | 258,096,837 | 17.55 | 19.55 | 92.05 |
| KLS1025 | *Hydrophis elegans* | 237,236,703 | 16.17 | 17.96 | 92.06 |
| KLS1027 | *Hydrophis elegans* | 241,492,305 | 16.48 | 18.42 | 92.09 |
| KLS1034 | *Hydrophis elegans* | 341,432,079 | 23.12 | 25.87 | 92.61 |
| KLS1121 | *Hydrophis elegans* | 821,569,381 | 54.77 | 61.41 | 93.46 |
| KLS0629 | *Hydrophis major* | 245,305,399 | 16.86 | 18.86 | 95.30 |
| KLS0630 | *Hydrophis major* | 224,462,436 | 15.43 | 17.26 | 95.20 |
| KLS0631 | *Hydrophis major* | 218,376,833 | 14.78 | 16.51 | 94.33 |
| KLS0647 | *Hydrophis major* | 173,722,269 | 11.96 | 13.33 | 94.70 |
| KLS1020 | *Hydrophis major* | 182,571,745 | 12.51 | 14.57 | 95.03 |
| FMNH249563 | *Hydrophis ornatus* | 174,414,840 | 11.83 | 13.23 | 90.92 |
| KLS1272 | *Hydrelaps darwinensis* | 266,993,222 | 14.47 | 16.31 | 85.44 |
| R36639 | *Hydrophis curtus* | 323,552,643 | 17.25 | 20.42 | 92.79 |
| KLS1291 | *Hydrophis kingii* | 1,070,617,691 | 63.78 | 75.24 | 95.99 |
| KLS1018 | *Hydrophis ocellatus* | 331,555,577 | 18.43 | 20.47 | 93.37 |
| KLS1292 | *Hydrophis peronii* | 319,639,177 | 17.54 | 19.53 | 93.29 |
| KLS0095 | *Hydrophis platurus* | 323,576,449 | 17.68 | 19.85 | 92.81 |
| KLS0026 | *Hydrophis schistosus* | 401,575,340 | 21.56 | 25.22 | 92.96 |
| LAHSc07 | *Hydrophis schistosus* | 355,348,815 | 19.37 | 22.55 | 93.01 |
| KLS1318 | *Hydrophis spiralis* | 578,123,505 | 35.78 | 40.42 | 95.09 |
| KLS0938 | *Hydrophis stokesii* | 318,032,773 | 17.12 | 19.07 | 93.40 |
| SD_mel1 | *Hydrophis cyanocinctus* | 89,389,145 | 3.78 | 4.46 | 78.97 |
| MT164 | *Hydrophis melanocephalus* | 55,978,594 | 1.99 | 2.23 | 60.61 |
| MT167 | *Hydrophis melanocephalus* | 56,083,949 | 1.94 | 2.25 | 59.10 |
| MT169 | *Hydrophis melanocephalus* | 99,983,384 | 4.54 | 5.34 | 82.04 |
| MT185 | *Hydrophis melanocephalus* | 86,845,505 | 3.79 | 4.25 | 78.16 |
| MT186 | *Hydrophis melanocephalus* | 87,081,255 | 3.23 | 3.81 | 73.95 |
| MT187 | *Hydrophis melanocephalus* | 85,670,500 | 3.47 | 3.97 | 71.23 |
| MW04737 | *Hydrophis melanocephalus* | 128,050,262 | 5.87 | 6.98 | 85.12 |
| MT04738 | *Hydrophis melanocephalus* | 122,529,175 | 5.86 | 6.87 | 86.02 |
| NN01 | *Naja naja* | 803,808,254 | 21.27 | 24.64 | 60.86 |
| NN02 | *Naja naja* | 536,952,687 | 29.85 | 34.56 | 60.34 |
| Notechis_scutatus | *Notechis scutatus* | 440,774,662 | 23.4 | 27.87 | 71.03 |
